# MELD: Mutation Effect Learning through Differential embeddings

**DOI:** 10.64898/2026.10.29.762727

**Authors:** Aliaa Alajji, Raihan Jashil, Raghvendra Mall, Mohamed Elshrif

**Author notes:** Work conducted during an internship at QCRI-HBKU.

## Abstract

We introduce MELD (Mutation Effect Learning through Differential Embeddings), a compact model for predicting the functional consequences of single nucleotide variants (SNVs) by training directly on the wild-type (WT) versus mutant differential. Interpreting SNVs is central to clinical genomics, yet experimental characterization cannot scale to millions of observed variants. Leading sequence-to-function models (Enformer, Borzoi, AlphaGenome) are trained on absolute genomic tracks and score variants post hoc through in-silico mutagenesis (ISM), so the WT-mutant difference never enters the training loss.

MELD instead places the logit difference Δ*z* = *z*_mut_ − *z*_wt_ inside the supervised objective. It combines a shared CNN-Transformer encoder, a mutant-specific mutation-mask boost, multiscale mutation-centered pooling, and a learned adaptive amplification factor *α*. On ClinVar (1.57M variants), MELD (1.9M parameters) reached an AUROC of 0.986 versus 0.800 for a same-data, same-backbone ISM baseline, with ablations confirming the contribution of each component. On a shared benchmark, it outperformed frozen foundation models with 3-1,300x more parameters (AUROC 0.967 vs. 0.924 for AlphaGenome and 0.936 for Nucleotide Transformer). With only 50, 000 training variants, it exceeded the ISM baseline trained on all 1.48M (AUROC 0.903 vs. 0.796) variants. On BRCA1 saturation genome editing, frozen MELD achieved a Spearman *ρ* of 0.42, above Borzoi (0.40), and ClinVar transfer raised it to 0.45. These results demonstrate that direct differential supervision is a compact, data-efficient alternative to post-hoc ISM scoring with substantially larger pretrained models.

## 1 Introduction

Determining how genetic variation produces functional change is a central problem in genomics and a prerequisite for interpreting the clinical significance of the human variome.^1–4^ Single nucleotide variants (SNVs) can alter gene regulation, chromatin accessibility, and transcriptional output, yet their effects are typically small, strongly context dependent, and difficult to measure at scale.^5–8^ Sequencing has made variant discovery routine, but exhaustive experimental characterization of variant function remains infeasible on cost and throughput grounds, leaving the large majority of observed variants functionally uncharacterized.^2, 9–12^

Sequence-to-function foundation models^7, 13–20^ have become the dominant computational response to close this gap. Methods such as Enformer,^19^ Borzoi,^20^ and AlphaGenome^7^ map long genomic windows, spanning tens of thousands to over a million base pairs, to thousands of epigenomic and transcriptional tracks, and they achieve state-of-the-art accuracy on absolute signal prediction. Variant effects are then obtained after training through in-silico mutagenesis (ISM) where the reference and alternate sequences are scored independently and their difference is taken as the predicted effect. A parallel family of self-supervised DNA language models produces general-purpose sequence representations that are transferred to variant scoring in the same post-hoc fashion, by comparing wild-type and mutant embeddings.^21–26^

This design carries a structural limitation. The training objective is defined over absolute signal, so the wild-type versus mutant difference that actually defines a variant effect never enters the loss optimization. Sensitivity to nucleotide-level perturbation is acquired only indirectly, as a byproduct of accurate absolute prediction. The documented consequences are attenuated response to low-effect variants, unreliable direction-of-effect estimates, and substantial inference cost from the repeated large-model forward passes that ISM requires.

Across these backbones, variant interpretation follows a common recipe: a pretrained or task-trained model produces a sequence representation, and the wild-type versus mutant differential is formed over a backbone that was never optimized to encode it. The families differ in what representation is available. Sequence-to-function models^7, 19^ expose predicted assay tracks, so we form the differential over those tracks, while DNA language models^22–24^ expose only learned trunk embeddings, so we use a light-weight prediction head (one output variable) over those. We hold the evaluation scaffold fixed across all backbones: a frozen encoder, a lightweight paired head over the wild-type and mutant representations, and identical training, split, and selection. Under this protocol the reported score is a supervised classifier probability (classification) or predicted effect score (regression), not an in-silico-mutagenesis contrast and not a zero-shot likelihood ratio. The representation type therefore varies by model class, which we state explicitly, while the head, data, and selection are matched.

Here we introduce MELD (Mutation Effect Learning through Differential Embeddings), a sequence-to-function model that places the wild-type versus mutant differential directly inside the supervised objective. A shared CNN-Transformer encoder processes matched reference and mutant windows, and the paired logit difference Δ*z* = *z*_mut_ − *z*_wt_ becomes the quantity the model is trained to predict on. A mutation mask marks the variant position and applies a learned boost to the mutant branch at that location, making the encoder sensitive to the altered site while retaining the surrounding sequence context. Wild-type and mutant representations are then pooled at a global scale and at increasingly local mutation-centered neighborhoods, so the differential is read out across multiple context scales rather than at a single resolution. A learned per-scale factor *α* provides context-dependent amplification of each scale-specific difference. Variant effect, thereby, becomes a quantity the model predicts directly rather than one recovered post hoc. In contrast to the frozen-transfer-based baselines, the representation itself is shaped to make wild-type versus mutant separation available, and we show this can be achieved with a compact encoder rather than a large pretrained foundation model. An overview of the classification and regression formulations is shown in Figure (1).

**Fig 1:**
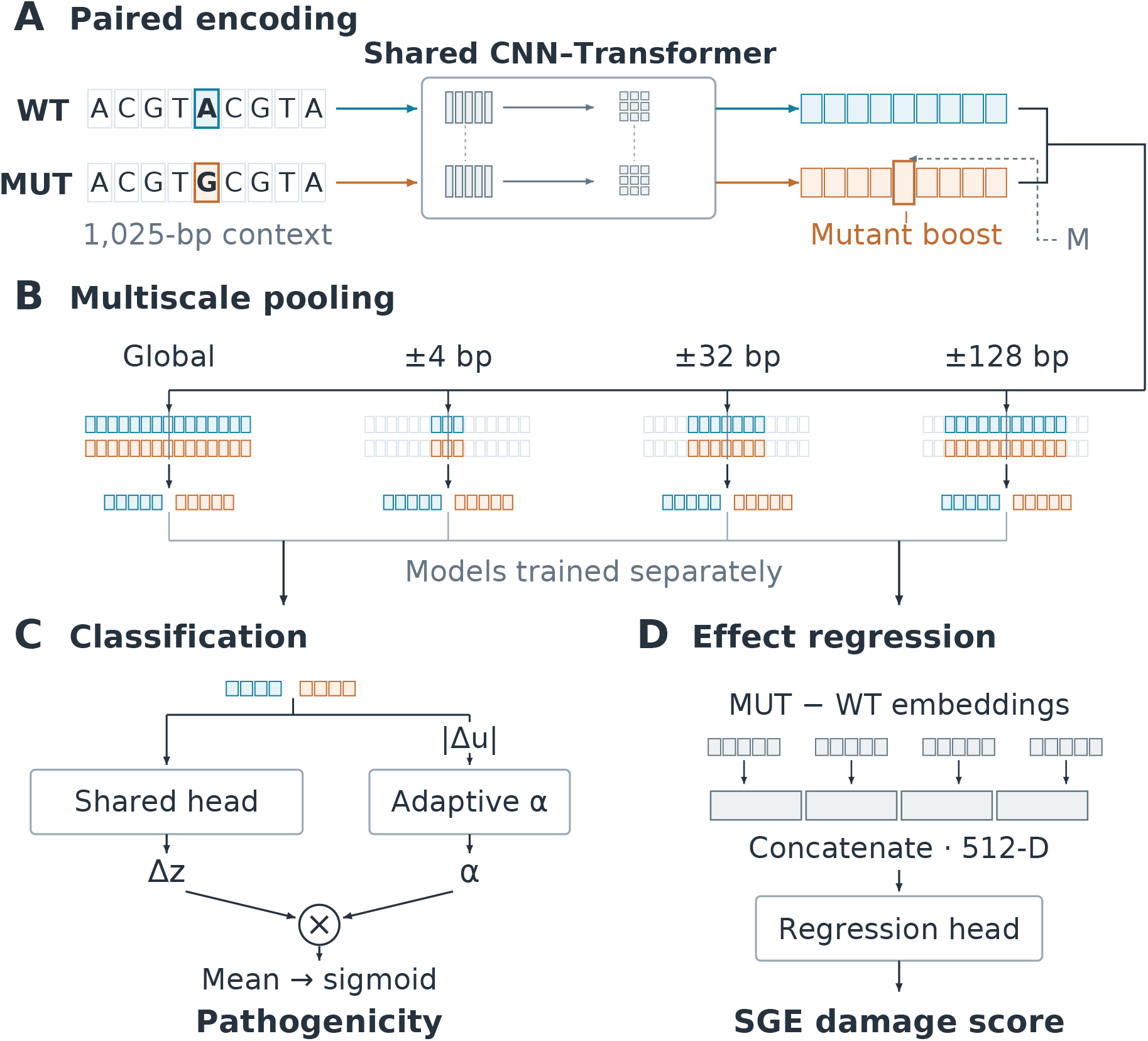
Overview of MELD for pathogenicity classification and quantitative effect prediction. **(A)** Paired wild-type (WT) and mutant (MUT) sequences are encoded using shared CNN-Transformer weights; mutation mask *M* guides mutant-specific modulation. **(B)** Global and mutation-centred pooling generate four pairs of 128-dimensional representations. **(C)** At each scale, a shared scalar head scores WT and MUT separately. Their MUT-minus-WT score difference is amplified by *α*, derived from the absolute embedding difference. Amplified differences are averaged across scales and passed through a sigmoid. **(D)** DeltaEmbedding regression concatenates the four MUT-minus-WT embedding differences into a 512-dimensional vector, which a regression head maps to a continuous saturation genome editing (SGE) damage score.

We evaluate MELD on ClinVar variant pathogenicity classification and on BRCA1 saturation genome editing (SGE) regression under matched data and backbone conditions. On the ClinVar held-out test set, MELD reaches an AUROC of 0.986, compared with 0.800 for a same-data, same-backbone ISM baseline that scores variants post-hoc, supporting the advantage of the paired MELD formulation over this baseline. With roughly 1.9 million parameters, MELD outperforms frozen pretrained foundation models with 3-1,300x more parameters when their representations are transferred to the same benchmark. The advantage holds across supervised-data regimes spanning ten thousand to more than a million training variants, and it extends from binary pathogenicity to continuous experimental effect. On BRCA1 SGE under a matched frozen-representation protocol, MELD attains a Spearman correlation (*ρ*) of 0.42, exceeding the strongest external encoder (Borzoi^20^), and end-to-end transfer from ClinVar pathogenicity predictor raises the *ρ* to 0.45. These results establish direct differential supervision as a compact and data-efficient framework for variant effect prediction.

## 2 Related Work

Computational variant effect prediction has been approached through two broad families of sequence models. The first family learns to predict experimental functional-genomics measurements directly from DNA, treating variant effect as a downstream consequence of an accurate sequence-to-function map. The second family pretrains large models on genomic sequence in a self-supervised fashion, producing general-purpose representations that transfer to many tasks. In both families, a variant is typically scored *post hoc*, by comparing model outputs or representations for the wild-type and mutant sequences after the backbone has been trained.

### 2.1 Sequence-to-function regression models

These models predict experimental coverage tracks directly from sequence. In their original formulations, variant effects are recovered through in-silico mutagenesis: the trained model is applied to the reference and alternate alleles and the resulting predicted tracks are contrasted, with no additional supervised head. We provide details about such methods below:

**Enformer:**^**19**^ Enformer pairs a convolutional tower with transformer layers, using self-attention to extend its effective receptive field well beyond earlier convolutional architectures.^19^ It takes an input window of roughly 200 kb and predicts thousands of human and mouse tracks, spanning CAGE, DNase/ATAC, and histone- and TF-ChIP, at 128 bp resolution, trained as a multitask regression against experimental coverage. By improving gene-expression prediction from sequence, it yields, as a downstream consequence, more accurate variant-effect predictions, obtained by contrasting model outputs for the reference and alternate alleles.

**Borzoi:**^**20**^ Borzoi extends this paradigm from summary expression values to full RNA-seq coverage profiles, predicting cell-type- and tissue-specific coverage directly from sequence.^20^ It adapts a convolutional and transformer backbone with a U-net-style upsampling path and processes a ∼524 kb context, substantially longer than Enformer’s, allowing it to score variant effects across transcription, splicing, and polyadenylation rather than a single expression summary. Borzoi is evaluated on quantitative trait loci and is competitive with and often surpasses models trained on individual regulatory functions.

**AlphaGenome:**^**7**^ AlphaGenome unifies and scales this direction further, taking up to 1 Mb of DNA as input and predicting thousands of functional-genomics tracks at single-base-pair resolution across many regulatory modalities, including gene expression, chromatin accessibility, histone modifications, transcription-factor binding, contact maps, and splicing.^7^ Its U-Net-style architecture downsamples the sequence through a convolutional encoder, exchanges information across positions with transformer layers, and upsamples back to base resolution before assay-specific out-put heads. It is trained in a supervised regime to predict experimental tracks and reports matching or exceeding the strongest external models on the large majority of its variant-effect benchmarks.

### 2.2 Self-supervised DNA language models

These models are pretrained with self-supervised objectives i.e. masked or autoregressive language modeling and have no task-specific supervision during pretraining; their representations transfer to downstream prediction either by fine-tuning or by training lightweight heads on frozen features.

**Nucleotide Transformer:**^**21**^ The Nucleotide Transformer (NT) family spans 50M to 2.5B parameters and is trained on progressively broader corpora, from the human reference genome to thousands of genetically diverse human genomes to genomes from hundreds of species, using non-overlapping 6-mer tokenization with a context of up to 1,000 tokens.^21^ After pretraining, its representations transfer to molecular-phenotype prediction by fine-tuning or by training light heads on frozen embeddings, with the largest multi-species variant generally strongest in the original benchmarking. Variant effects are derived by comparing representations for wild-type and mutant sequences.

**DNABERT-2:**^**22**^ DNABERT-2 is a compact (∼117M-parameter) BERT-style encoder that replaces the overlapping *k*-mer tokenization of the original DNABERT with byte-pair encoding and swaps learned positional embeddings for Attention with Linear Biases, alongside efficiency-oriented architectural changes.^22^ Pretrained on a large multi-species corpus, it reports performance comparable to or better than the 2.5B-parameter Nucleotide Transformer^21^ on a multi-task genome-understanding benchmark introduced alongside it, while using roughly 21x fewer parameters and approximately 92x less GPU time in pretraining. Its size makes it a practical backbone for genomic sequence-classification tasks under constrained compute.

**Caduceus:**^**23**^ Caduceus is a family of bi-directional, reverse-complement (RC) equivariant long-range DNA language models built on the Mamba state-space block.^23^ It extends Mamba to a Bi-Mamba component for bi-directional context and a MambaDNA block that enforces RC equivariance, and stacks these layers into a bidirectional trunk that can be pretrained on chromosome-length windows with only a few million parameters. Pretrained by masked language modeling on the human reference genome, it reports that on a challenging long-range variant-effect task it exceeds transformer baselines roughly an order of magnitude larger that do not exploit bi-directionality or strand symmetry. As with the other language models, variant effects are read out post hoc by contrasting learned representations for the wild-type and mutant sequences.

**HyenaDNA:**^**24**^ HyenaDNA replaces attention with the implicit long-convolution Hyena operator, giving single-nucleotide-resolution modeling with full global context at every layer and sub-quadratic scaling in sequence length.^24^ This lets it reach context lengths of up to one million tokens - an up to 500× increase over prior dense-attention genomic models - while training substantially faster. Pretrained autoregressively by next-token prediction on the human reference genome with far fewer parameters and less data than earlier foundation models, it reaches state-of-the-art on many of the Nucleotide Transformer and GenomicBenchmarks datasets. Variant effects are again obtained post hoc, by comparing representations for the reference and alternate sequences.

### 2.3 Positioning

For the frozen-representation comparison, sequence-to-function models provide predicted assay tracks, whereas DNA language models provide trunk embeddings. Wild-type and mutant sequences are processed independently using a fixed pretrained backbone, and differences between their representations are supplied to a task-specific downstream head. The reported outputs are supervised pathogenicity probabilities for classification or continuous mutation-effect scores for regression, rather than unmodified native zero-shot scores.

In addition to frozen-representation baselines, we evaluate full end-to-end fine-tuning of DNABERT-2,^22^ Caduceus^23^ and HyenaDNA^24^ for ClinVar pathogenicity classification, updating the pretrained backbones as well as their prediction heads. This extends the comparison beyond heads trained on fixed features to models whose representations are adapted to the downstream task.

MELD combines a compact shared CNN–Transformer encoder with mutation-aware modulation and multiscale pooling. Classification uses adaptively amplified scalar differences, whereas quantitative regression uses concatenated embedding differences and a separate regression readout. These formulations allow efficient paired mutation-effect prediction without requiring a large pretrained foundation-model backbone.

## 3 Methods

### 3.1 Problem formulation

Each example consists of a wild-type DNA sequence window *S*_*wt*_, a matched mutant sequence window *S*_*mut*_, and a variant-effect label *y*. For pathogenicity classification, *y* ∈ {0, 1}, where 0 denotes benign and 1 denotes pathogenic; for quantitative mutation-effect regression, *y* ∈ R is a continuous experimental effect score. The objective is to predict the effect of the mutation rather than the absolute function of either sequence alone.

Conceptually, the mutation-effect signal is represented as the difference between mutant and wild type logits:

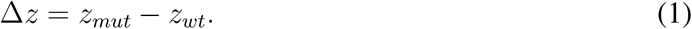

The paired differential score can be expressed in simplified form as

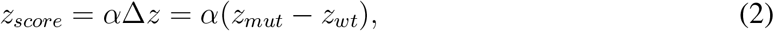

followed by a sigmoid probability:

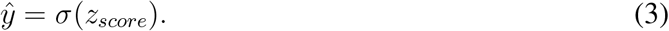

In our MELD framework, this differential score is computed separately across multiple mutation-context scales using scale-specific adaptive amplification factors, and the resulting amplified logits are averaged to obtain the final score. Classification uses an amplified scalar differential, whereas quantitative regression uses the DeltaEmbedding readout, which concatenates multiscale differences between pooled WT and mutant representations and maps them through a nonlinear regression head with a final linear output, detailed in Section 4.7.

This formulation places the WT-mutant differential directly inside the supervised training objective. This differs from post-hoc variant scoring, where a model is first trained to predict an absolute sequence output and mutation effects are later estimated by comparing predictions for reference and alternative alleles.

### 3.2 Shared WT/mutant encoder

Wild-type and mutant sequences are encoded using the same shared-weight encoder *f*_*θ*_:

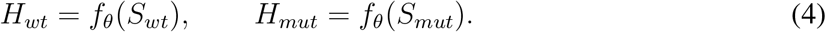

The encoder is shared between the two branches, so WT and mutant representations are produced in the same learned feature space. The active implementation uses a compact CNN-Transformer encoder composed of token and positional embeddings, five residual CNN blocks, one Transformer encoder layer, and LayerNorm:

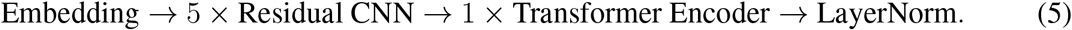

This compact backbone is used to test the paired differential learning objective under a controlled model capacity. The primary contribution is the paired WT/mutant objective and differential scoring mechanism, not the use of a large pretrained sequence model.

### 3.3 Mutation-mask boost

A binary mutation mask *M* ∈ {0, 1}^*L*^ marks the position or span affected by the variant. For SNVs, a single base is marked. For indels, the affected span is defined as max(1, |REF|, |ALT|) positions beginning at the variant position.

The mutant hidden states are modulated using a learned non-negative parameter *β*:

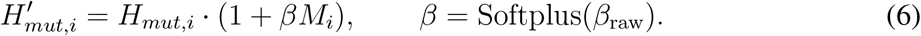

Only the mutant branch is boosted. The wild-type hidden states are not modified by the mutation-mask boost. This mechanism encourages the model to emphasize mutation-affected positions while retaining information from the full encoded sequence.

### 3.4 Multi-scale mutation-context pooling

After encoding, the model pools WT and mutant representations at multiple context scales around the mutation. The active radii are:

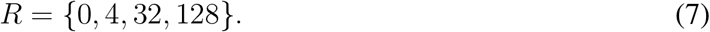

In the current implementation, radius *r* = 0 corresponds to global masked mean pooling over the full valid input window, excluding padding positions. Radii *r >* 0 correspond to mutation-neighborhood masked mean pooling, where the mutation mask is expanded to include positions within radius *r* and intersected with the valid-token mask.

For each radius *r*, pooled representations are computed:

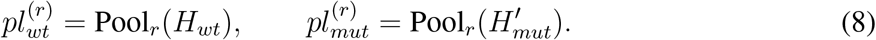

A shared LayerNorm-plus-linear head is applied independently to the pooled representations of both alleles at each scale:

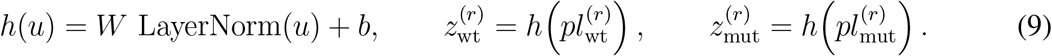

The same head parameters are shared across alleles and pooling scales. The scale-specific mutation signal is:

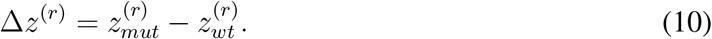

The use of multiple pooling scales allows the model to combine global sequence context with increasingly localized mutation-centered summaries.

### 3.5 Adaptive logit amplification

For each pooling scale *r*, the model computes an adaptive amplification factor from the absolute difference between pooled mutant and WT representations:

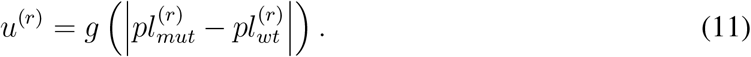

Here, *g*(·) is shared across pooling scales and consists of LayerNorm, a 128-unit linear layer, GELU activation, dropout 0.15, and a scalar linear output. The raw nonnegative amplification contribution is produced using Softplus:

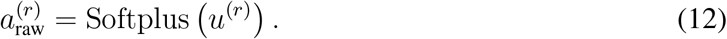

The learned base amplification factor is:

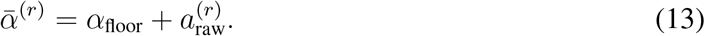

The epoch-dependent effective amplification factor is:

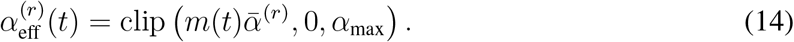

Here, *α*_floor_ is the minimum amplification offset, *α*_max_ is the maximum allowed amplification, and *m*(*t*) is the epoch-dependent alpha curriculum multiplier. In MELD configuration, *α*_floor_ = 1.0 and *α*_max_ = 48.0. We set *α*_floor_ = 1 so that, at full-strength evaluation, adaptive amplification cannot attenuate the underlying WT–mutant logit differential. The upper bound *α*_max_ = 48 prevents unbounded amplification and provides a finite numerical cap. The sensitivity analysis additionally evaluated smaller amplification bounds, showing that performance was relatively insensitive to the precise upper-bound value.

The scale-specific amplified mutation logit is:

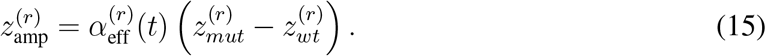

The final logit is averaged across pooling scales:

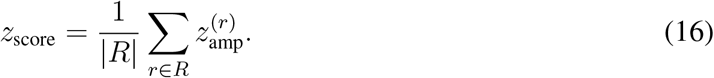

### 3.6 Alpha curriculum

During early training, full amplification can be unstable because WT and mutant representations are not yet well formed. The implementation therefore uses an alpha curriculum that gradually increases the effective amplification strength during the first training epochs.

Let *m*(*t*) denote the curriculum multiplier at the one-based training epoch *t*:

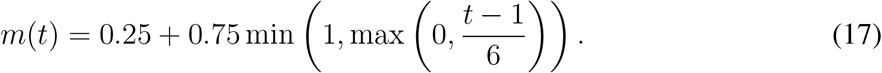

Thus, *m*(1) = 0.25, the multiplier increases linearly by 0.125 per epoch, and reaches 1.0 at epoch 7. This means that the learned alpha module is active throughout training, but its effective strength is reduced at the start and progressively increased as the encoder representations become more stable. The multiplier is applied once to the sum of the amplification floor and learned nonnegative contribution before clipping. From epoch 7 onward, the model uses the full learned amplification factor, subject to the configured clipping range.

### 3.7 Training objective

For binary classification, the primary loss is focal binary cross-entropy with automatic positive-class weighting. Let *z*_score,*i*_ denote the final paired-model logit for variant *i*, and let *y*_*i*_ ∈ {0, 1} denote its binary label. The predicted probability is:

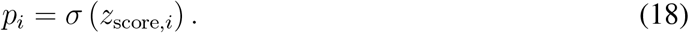

For each example, define:

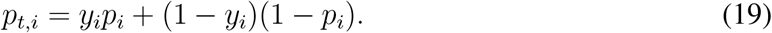

The positive-class-weighted binary cross-entropy term is:

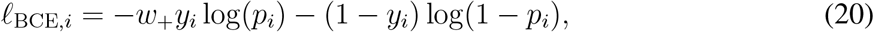

where *w*_+_ = *N*_−_*/N*_+_ is computed from the training-label distribution. The focal classification loss is then defined as:

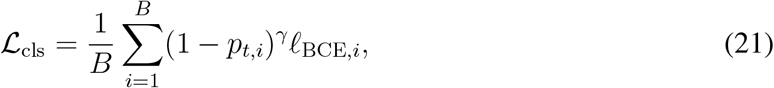

where *B* is the minibatch size and *γ* is the focal-loss parameter and has been shown to be effective for imbalanced classification tasks.^27, 28^

In addition to the classification loss, the model uses an auxiliary delta regularization term on the WT/mutant logit difference. For each variant, Δ*z*_*i*_ is computed from the unamplified, scale-averaged WT and mutant logits:

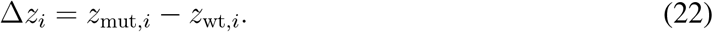

Let P = {*i* : *y*_*i*_ = 1} denote pathogenic variants and N = {*i* : *y*_*i*_ = 0} denote benign variants. The delta regularization term is:

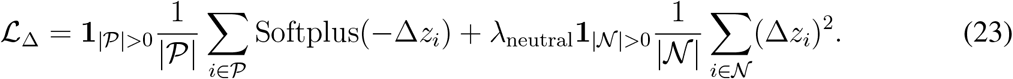

The indicator terms skip the corresponding component when a minibatch contains no examples from that class. The first term encourages pathogenic variants to satisfy *z*_mut_ *> z*_wt_, so that the mutant sequence receives a higher pathogenicity logit than the matched WT sequence. The second term weakly regularizes benign variants toward Δ*z* ≈ 0, rather than forcing benign mutants to satisfy *z*_mut_ *< z*_wt_. This reflects the expectation that benign variants should have little predicted effect, rather than necessarily having a protective effect.

The total training objective is:

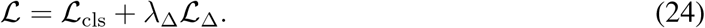

In the standard classification configuration, *λ*_Δ_ = 0.05 and *λ*_neutral_ = 0.1; both regularization weights were included in the hyperparameter sensitivity analysis. The focal-loss parameter *γ* was fixed at 1.0, while positive-class weighting was computed automatically from the training-label distribution as *N*_−_*/N*_+_.

### 3.8 Same-backbone post-hoc ISM baseline

To test whether direct paired WT/mutant training improves over post-hoc variant scoring, we implemented a same-data, same-backbone ISM-style baseline. This baseline uses the same compact CNN-Transformer encoder family as MELD, but trains it as an absolute single-sequence classifier rather than as a paired WT/mutant model.

For each variant *i*, let 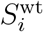 and 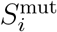 denote the wild-type and mutant sequence windows, and let *y*_*i*_ ∈ {0, 1} denote the variant label, where 0 indicates benign and 1 is pathogenic. The baseline converts each paired variant row into two single-sequence training examples with labels 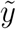:

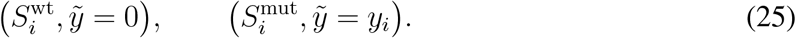

Thus, benign variants contribute two negative examples, one from the WT sequence and one from the mutant sequence. Pathogenic variants contribute a negative WT example and a positive mutant example. The baseline is therefore trained on individual sequences and does not receive the WT/mutant pair jointly during training.

At evaluation time, the trained single-sequence classifier is applied independently to the WT and mutant sequences. If *f*_*θ*_(*S*) denotes the scalar single-sequence logit, the variant pathogenicity probability is:

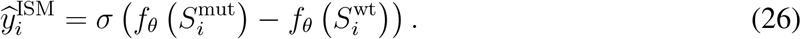

The mutant-minus-WT sign is fixed before evaluation. Checkpoints are selected using validation area under precision-recall curve (AUPRC), and the classification threshold is selected by maximizing validation F1. This provides a controlled same-data, same-backbone comparison between direct paired supervision and post-hoc differential scoring.

## 4 Experimental Setup

### 4.1 Primary benchmark: ClinVar standard 1kb

The primary benchmark was constructed from ClinVar^29^ using nominal 1,025-bp variant-centred DNA windows. Each row contains a paired wild-type and mutant DNA window, variant metadata, a binary pathogenicity label, and a preserved train/validation/test split. The split and label distribution are shown in Table 1. The label mapping is: 0 = benign, 1 = pathogenic. The dataset contains 1,572,808 paired variant rows.

**Table 1:** ClinVar standard 1 kb dataset split and label distribution.

| **Split** | **Total rows** | **Benign** | **Pathogenic** |
|---|---|---|---|
| Train | 1,482,739 | 1,198,246 | 284,493 |
| Validation | 35,556 | 29,127 | 6,429 |
| Test | 54,513 | 46,120 | 8,393 |
| Total | 1,572,808 | 1,273,493 | 299,315 |

The sequence windows are constructed using the hg38 reference genome. The provided train / validation / test split is preserved exactly across all main models, ablations, and same-backbone baseline experiments. No reshuffling or resplitting is performed.

### 4.2 ClinVar dataset description

Before final reporting, the ClinVar standard dataset was audited to avoid any split leakage and data-quality issues. The split is chromosome-disjoint, and no cross-split overlap was found at the variant, WT/mutant-pair, or exact sequence-window level. The dataset is ClinVar-only, so no source-label confounding warning was raised.

We performed an audit that identified a small number of duplicated WT/mutant pairs and duplicated sequence windows. These duplicates were documented and retained because they did not introduce cross-split leakage, and the same audited dataset and fixed split were used consistently across all models and baselines. The audit also identified multi-allelic REF positions, which were retained and documented for transparency.

In addition to exact split-leakage checks, we performed a sequence-similarity analysis using a MinHash/LSH near-duplicate screen, recording for each validation or test sequence the nearest approximate match in the training set. No validation or test sequence exceeds 0.90 sequence-similarity to the training set; the only nonzero hit was a single mutant validation sequence at 0.875, below the near-duplicate threshold. Full per-split results are reported in Supplementary Table S1. This supports that the held-out performance is not explained by trivial sequence-window leakage.

### 4.3 BRCA1 saturation genome editing dataset

For quantitative mutation-effect evaluation, we used the BRCA1 saturation genome editing (SGE) dataset of Findlay et al.,^30^ accessed through MaveDB^31^ (URN: urn:mavedb:00000097-0-2). The deposited score set contains 3,893 variants, of which 2,859 unique SNVs have usable GRCh38 genomic mappings and pass reference-allele and sequence-construction checks. The regression target was defined as the sign-aligned damage score, obtained by negating the deposited raw SGE score so that larger values indicate greater functional impairment.

To reduce leakage from overlapping local sequence contexts, the 2,859 variants were partitioned using 12 nonoverlapping sequence-window components. The resulting fixed split contains 1,995 training variants, 398 validation variants, and 466 held-out test variants as depicted in Fig. 2. The largest component contains 343 variants (12.0% of the dataset), the minimum gap between sequence windows assigned to different splits is 393 bp, and no stable variant identifier or reference sequence window overlaps across splits. The same fixed split and continuous target are used for the BRCA1 regression comparisons reported here.

**Fig 2:**
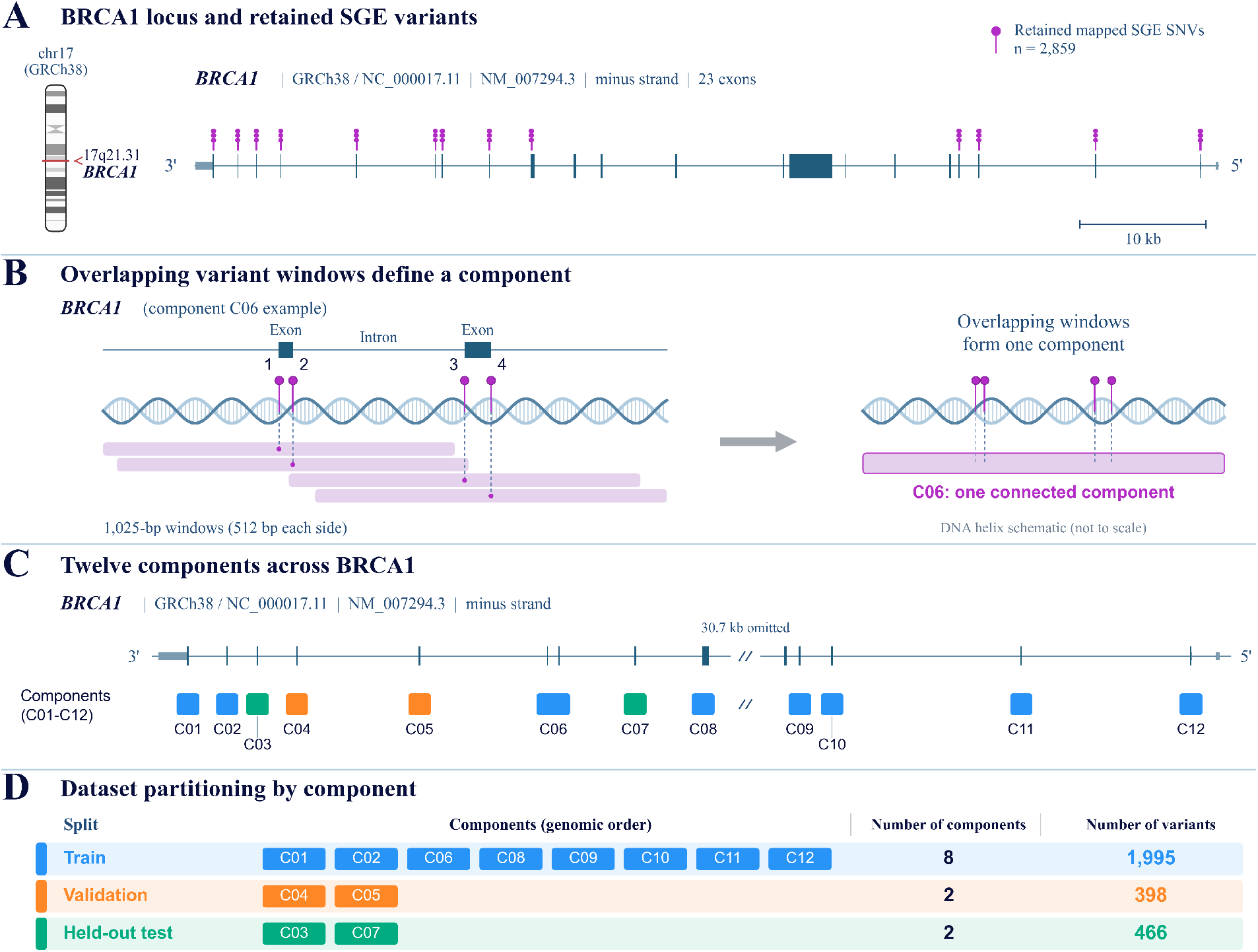
BRCA1 saturation genome editing variants partitioned by overlapping sequence context. **(A)** Genomic distribution of the retained BRCA1 SGE variants. **(B)** Example overlap-connected component illustrating how overlapping 1,025-bp sequence windows are grouped for split assignment. **(C)** The 12 sequence-window components used to construct the grouped train, validation, and test partitions. **(D)** Summary of the fixed split containing 1,995 training, 398 validation, and 466 held-out test variants.

### 4.4 Preprocessing and input construction

For the ClinVar classification task, each example contains an aligned wild-type sequence *S*_wt_, mutant sequence *S*_mut_, binary pathogenicity label *y*, and binary mutation mask *M*. The WT and mutant windows are processed as a paired input and retain the original ClinVar train, validation, and test assignments.

When genomic variant metadata are available, the mutation mask marks max(1, |REF|, |ALT|) positions beginning at the variant position. Otherwise, the affected positions are inferred from differences between the WT and mutant sequences. Valid-token attention masks are used to exclude padding positions during pooling.

### 4.5 Data augmentation

During standard training, reverse-complement augmentation is applied with probability 0.5. A single Bernoulli draw is made for each minibatch, and the WT sequences, mutant sequences, and mutation masks are reverse-complemented jointly to preserve their alignment.

The standard configuration also applies positional-shift augmentation. One shared shift is sampled uniformly from [−8, 8] for each minibatch and applied consistently to the WT sequences, mutant sequences, attention masks, and mutation masks. Setting the maximum shift to zero disables positional-shift augmentation.

During evaluation, reverse-complement test-time augmentation is applied by computing logits for the original and reverse-complement inputs and averaging the two logits before applying the sigmoid function. Reverse-complement consistency metrics are also reported during evaluation, but reverse-complement consistency loss is not part of the standard training objective.

### 4.6 Classification model configuration

MELD uses the shared CNN-Transformer encoder described in Section 3, with 1,917,059 trainable parameters, comprising token and positional embeddings, five residual dilated CNN blocks, one Transformer encoder layer, and a final LayerNorm (Eq. 5). The WT and mutant windows are processed by the same shared encoder and prediction head. Representations are pooled globally and within mutation-centered neighborhoods using the pooling radii *R* ∈ {0, 4, 32, 128}, where *R* = 0 denotes masked mean pooling over the complete valid sequence and radii 4, 32, and 128 denote masked mean pooling over mutation-centered neighborhoods. The per-scale effective amplification factor (Eq. 14) uses *α*_floor_ = 1 and *α*_max_ = 48; the final paired-model logit is the scale-averaged amplified differential (Eq. 16), mapped to a pathogenicity probability by the sigmoid (Eq. 3).

The standard training objective combines positive-class-weighted focal binary cross-entropy with an auxiliary WT/mutant delta regularizer. The standard configuration uses focal-loss parameter *γ* = 1, automatic positive-class weighting, delta-regularizer weight *λ*_Δ_ = 0.05, and neutral-variant regularization weight *λ*_neutral_ = 0.1.

### 4.7 Quantitative regression formulations

For quantitative mutation-effect prediction on the BRCA1 saturation genome editing dataset, we evaluated MELD-DeltaEmbedding and a same-backbone ISM regression baseline. Both use the compact 5-CNN/1-Transformer encoder family and the SmoothL1 regression loss.

#### MELD-ΔEmbedding

For each pooling scale *r* ∈ {0, 4, 32, 128}, we compute the difference between the pooled mutant and wild-type representations,

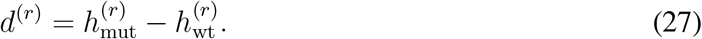

The four 128-dimensional differential representations are concatenated,

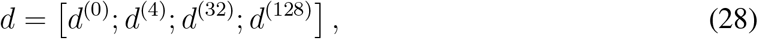

and mapped to a continuous mutation-effect prediction using a small regression head consisting of LayerNorm, a 512-to-128 linear layer, GELU activation, dropout of 0.15, and a final scalar linear layer. No sigmoid or adaptive-alpha scaling is applied in this regression head. The model contains 1,966,594 trainable parameters.

#### MELD initialization and ClinVar transfer

For the random-initialization condition, all MELD parameters were initialized randomly and the model was trained directly on the 1,995 BRCA1 training variants, with checkpoint selection performed using Spearman correlation on the 398-variant BRCA1 validation set. For the strict ClinVar-transfer condition, MELD was first pretrained on the ClinVar pathogenicity task after excluding BRCA1-related training examples whose variant alleles or local sequence contexts overlapped the BRCA1 regression benchmark. The resulting pretrained weights were then used to initialize MELD before fine-tuning on the same 1,995 BRCA1 training variants. Both conditions used the same 398-variant BRCA1 validation set for checkpoint selection and the same 466-variant held-out test set for final evaluation.

#### Same-backbone ISM regression baseline

For the controlled post-hoc baseline, the same encoder family is trained as an absolute single-sequence regressor. For each variant, the wild-type sequence is assigned target 0, whereas the mutant sequence is assigned the continuous experimental mutation-effect target. At evaluation, the predicted mutation effect is obtained as

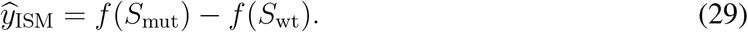

This baseline contains 1,900,161 trainable parameters. MELD-DeltaEmbedding forms the WT/mutant differential in representation space before regression, whereas the ISM baseline forms the difference after independent sequence scoring.

### 4.8 External baselines for quantitative-effect regression

To place MELD’s quantitative-effect predictions in the context of pretrained genomic sequence models, we evaluated frozen external encoders using the same BRCA1 grouped train, validation, and held-out test split. The representation over which the wild-type versus mutant differential is formed depends on the model class. For the sequence-to-function models (Enformer,^19^ Borzoi,^20^ AlphaGenome^7^) we use their native predicted assay tracks, whereas for the four DNA language models (Nucleotide Transformer,^21^ DNABERT-2,^22^ Caduceus,^23^ HyenaDNA^24^) we use frozen trunk embeddings. In every case the pretrained encoder parameters remained frozen, and WT and mutant sequences were encoded independently.

The primary external-baseline formulation uses the differential in representation space. Given frozen WT and mutant representations, *h*_WT_ and *h*_MUT_, we compute

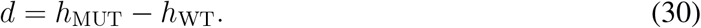

The resulting differential representation is passed through a lightweight regression head consisting of layer normalization, a linear projection to 128 dimensions, GELU activation, dropout of 0.15, and a final linear layer producing an unrestricted continuous prediction. No sigmoid, ReLU, or other output-bounding activation is applied.

All downstream heads are trained using the BRCA1 training set only, with checkpoint and representation-layer selection performed using validation Spearman correlation. We used five different random seeds for each configuration. Encoder architecture, representation extraction, pooling strategy, and downstream-head configuration are fixed before evaluation on the held-out set.

### 4.9 Precomputed genome-wide baselines

To compare MELD with established genome-wide variant scores, we evaluated four precomputed predictors: CADD v1.7,^32^ GPN-MSA,^33^ and the 100-way vertebrate phyloP^34^ and phastCons^35^ conservation scores. None of these scores was trained or tuned on our data. Each was taken from its published hg38 release and joined to the ClinVar validation and test variants by genomic coordinate. CADD (PHRED score) and GPN-MSA were matched on chromosome, position, reference and alternate allele, and phyloP and phastCons on position. GPN-MSA scores were negated so that higher values indicate greater predicted pathogenicity; the other three scores already follow this direction.

The CADD and GPN-MSA score files cover single-nucleotide variants only, so this comparison is restricted to SNVs scored by both: 48,795 of the 48,835 SNVs in the full held-out test set and 17,908 of the 17,918 SNVs in the shared-subset test set. MELD was evaluated on the same variants using its existing per-variant test predictions. AUROC and AUPRC were computed from the raw scores. F1 and MCC used the threshold that maximized F1 on the corresponding validation SNVs, applied once to the test set, as for all other models. Calibration metrics are not reported because these scores are not probabilities. The precomputed scores are fixed lookups with no seed variability; MELD values are five-seed mean ± standard deviation.

### 4.10 Architecture capacity sweep

The MELD configuration with five residual CNN blocks and one Transformer encoder layer is used as the initial standard architecture. To determine whether model depth improves mutation-effect prediction, we compare encoder variants with different numbers of residual CNN blocks and Transformer encoder layers while fixing the paired WT/mutant training objective, data split, and training setup. We also include CNN-only and Transformer-only encoder ablations to assess whether local convolutional feature extraction or Transformer-based contextualization contributes more strongly to performance.

The 5CNN+2T configuration isolates the effect of adding Transformer depth without increasing CNN depth. The 7CNN+2T configuration tests a moderate increase in both local CNN capacity and Transformer depth. The 7CNN+4T configuration tests whether deeper Transformer capacity improves contextual interaction modeling. The CNN-only 5CNN model removes Transformer layers entirely to test whether local convolutional sequence features are sufficient, while the Transformer-only 2T model removes CNN blocks to test whether self-attention alone can recover the relevant local sequence patterns.

### 4.11 Hyperparameter sensitivity analysis

We performed a staged, grouped hyperparameter sensitivity analysis using the compact 5CNN+1T architecture. A WanDB sweep using Bayesian optimization^27,36^ was performed over 36 unique configurations. The configurations independently varied optimizer settings, delta-regularizer weights, alpha formulation and upper bound, reverse-complement and positional-shift augmentation, or dropout.

Each configuration started from the same compact base configuration, and later sensitivity groups did not inherit the best settings from earlier groups. Checkpoints were selected using validation AUPRC, and classification thresholds were selected by maximizing validation F1. The held-out test set was evaluated only after validation-based checkpoint and threshold selection.

### 4.12 Baselines and ablations

To isolate the contribution of each modeling component, we compare MELD against the following baseline and ablation configurations:

- **Same-backbone ISM-style baseline:** the post-hoc differential baseline defined in Section 3.8, trained as an absolute single-sequence classifier over the same compact CNN– Transformer encoder family.
- **No-alpha ablation:** fixes the amplification factor to *α* = 1, reducing the prediction score to the unamplified paired WT/mutant differential.
- **No-mutation-boost ablation:** disables the learned mutation-mask boost applied to the mutant hidden states.
- **Single-scale ablation:** disables mutation-neighborhood pooling and uses only global valid-sequence pooling.
- **No-RC ablation:** disables reverse-complement augmentation during training and reverse-complement test-time augmentation during evaluation. Positional-shift augmentation remains enabled.
- **No-delta-regularizer ablation:** sets the auxiliary WT/mutant delta-regularizer weight to zero.
- **256-bp context ablation:** reduces the input sequence context from 1 kb to 256 bp while retaining the paired model formulation.
- **Residual-alpha variants:** replace the current amplification formulation with the residual-alpha formulation using *α*_max_ = 8 or *α*_max_ = 16.

The primary comparison is MELD versus the same-data, same-backbone ISM-style baseline. This controlled comparison tests whether placing the WT-mutant differential directly inside the supervised objective improves performance over post-hoc differential scoring.

### 4.13 Data-efficiency and training-set scaling

To evaluate how the paired MELD formulation behaves as the amount of supervised training data changes, we performed a controlled data-efficiency comparison against the same-backbone ISM-style baseline. Starting from the fixed ClinVar training split of 1,482,739 variants, we constructed deterministic nested training subsets containing 10,000, 50,000, 200,000, and 500,000 variants, together with the complete training set. The original validation set of 35,556 variants and held-out test set of 54,513 variants were kept unchanged for every training-size condition. The full-data condition was retrained under the same update-controlled protocol used for all learning-curve conditions and is therefore treated as a separate controlled retraining rather than a reuse of the primary MELD checkpoints.

The training subsets preserve the original ClinVar labels and are nested such that every smaller subset is contained within the next larger subset; the resulting training-set compositions are reported in Supplementary Table S2.

MELD and the same-backbone ISM baseline were trained independently at each data scale using five random seeds. To make optimization effort comparable across training-set sizes, training was controlled by optimizer updates rather than by a fixed number of epochs. The effective batch size was 128, with a maximum of 150,000 optimizer updates. Validation was performed every 1,500 updates, with a minimum of 6,000 updates before early stopping and a patience of 10 validation events, corresponding to 15,000 updates without improvement.

Model checkpoints were selected using validation AUPRC. For threshold-dependent classification metrics, the decision threshold was selected by maximizing validation F1 and was then applied unchanged to the held-out test set. The same training subsets, validation set, test set, and five seeds were used for MELD and ISM.

### 4.14 Evaluation metrics and selection protocol

We evaluate binary pathogenicity prediction using AUROC, AUPRC, F1, Matthews correlation coefficient (MCC), balanced accuracy, accuracy, precision, recall, specificity, false-positive rate (FPR), Brier score, and expected calibration error (ECE).

AUROC and AUPRC are computed from continuous predicted probabilities and are independent of the classification threshold. F1, MCC, balanced accuracy, accuracy, precision, recall, specificity, and FPR are computed using a classification threshold selected on the validation set by maximizing validation F1 and then applied once to the held-out test set. These metrics are widely used for binary and imbalanced data classification as illustrated in.^27, 28, 37^

The optimal model configuration is selected using validation AUPRC. Validation data are used only for configuration selection, threshold selection, early stopping, and training monitoring. The held-out test split is never used for model checkpoint, hyperparameter, or threshold selection.

Unless explicitly identified as a single-run sensitivity experiment, repeated-seed results use 5 different random seeds and are reported as mean ± standard deviation. The hyperparameter sensitivity analysis uses fixed seed of 42 and is reported separately from the five-seed main-model and ablation results.

For quantitative BRCA1 mutation-effect regression, performance is evaluated using Pearson correlation, Spearman rank correlation, root mean squared error (RMSE), mean absolute error (MAE), and coefficient of determination (*R*^2^). Regression checkpoints are selected using validation Spearman correlation. Regression outputs are continuous and unrestricted real-valued.

## 5 Results

### 5.1 Base model and architecture capacity

The MELD model was evaluated on the held-out ClinVar standard test split using 1,025-bp variant-centered sequence windows. This reference architecture contains 1,917,059 trainable parameters and reached a peak GPU memory usage of 10.22 GB during training.

The reference MELD model reaches AUROC 0.9859 ± 0.0007 and AUPRC 0.9342 ± 0.0052, with strong balanced performance (F1 0.8598 ± 0.0047, MCC 0.8358 ± 0.0050) and calibration (Brier 0.0472 ± 0.0009, ECE 0.0711 ± 0.0014). Full held-out metrics, including balanced accuracy, accuracy, precision, recall, specificity, and false-positive rate, are reported in Supplementary Table S3. These results show that a compact shared CNN-Transformer model trained on paired WT/mutant windows can achieve high held-out pathogenicity prediction performance without requiring a large pretrained foundation-model backbone, achieving both high ranking performance (AUROC, AUPRC) and strong thresholded classification (F1, MCC). This MELD model is therefore retained as a parameter-efficient reference architecture, while the architecture-capacity analysis evaluates whether increasing encoder depth improves performance.

We evaluated whether increasing encoder depth improves mutation-effect prediction beyond the compact MELD architecture. All architecture variants use the same paired WT/mutant objective, ClinVar standard split, input representation, and training protocol. The sweep varies the number of residual CNN blocks and Transformer encoder layers to assess whether additional local feature-extraction capacity or contextual modeling improves performance.

Across the architecture sweep, MELD achieved the strongest mean performance across all reported predictive and calibration metrics (Supplementary Table S4). From Fig. 3D, we observe that the 5CNN+2T model achieved nearly identical AUPRC to the 5CNN+1T architecture despite using an additional Transformer layer and ≈ 10% more parameters. Increasing both convolutional and transformer depth further reduced mean performance and increased variability across seeds, indicating that additional encoder capacity may lead to overfitting and does not consistently benefit this task.

**Fig 3:**
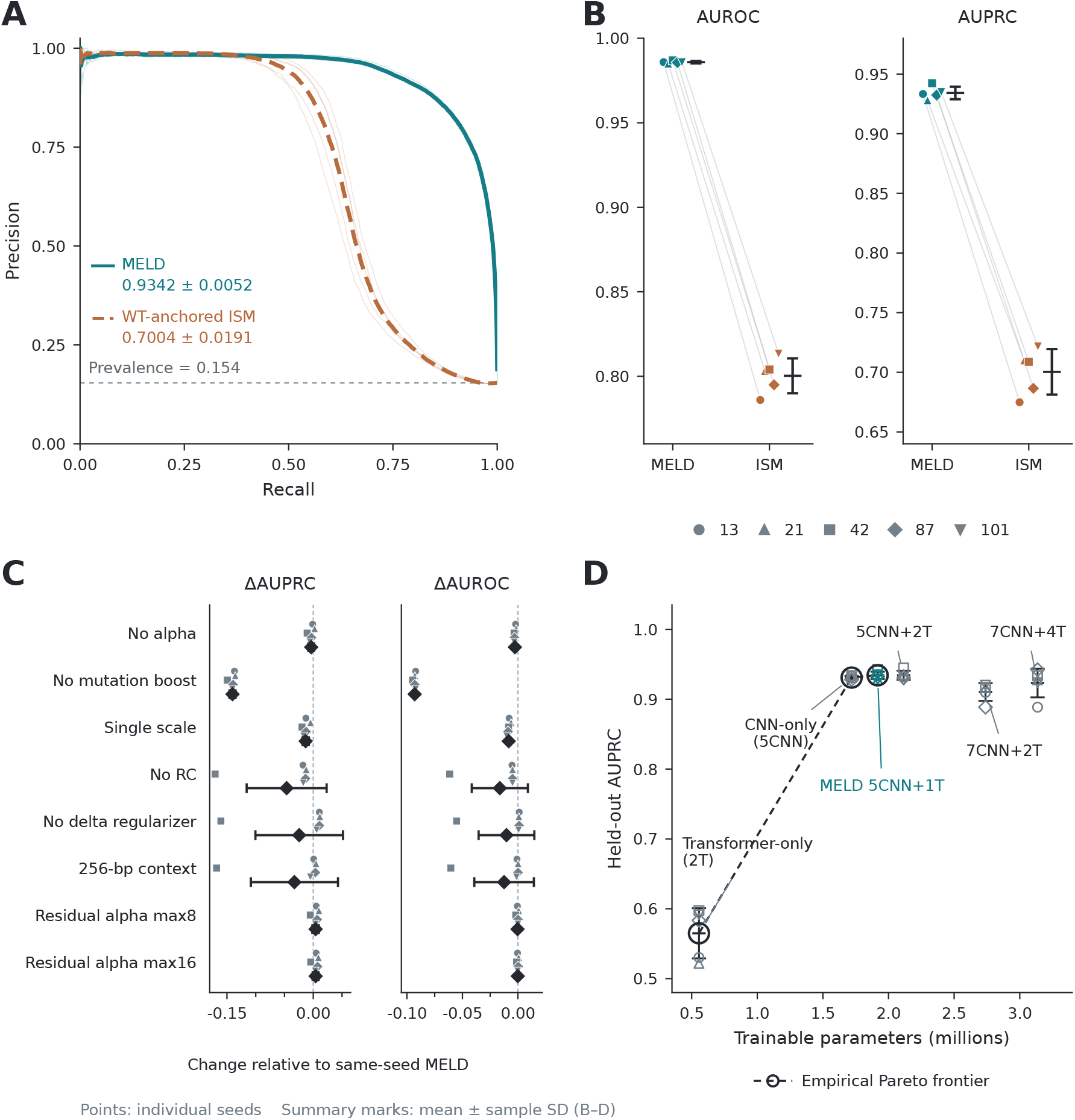
Primary ClinVar performance and model-design analysis. **(A)** Precision–recall curves for MELD and the same-backbone ISM baseline across five seeds; bold curves denote the seed-averaged interpolation and the dashed horizontal line indicates test-set pathogenic prevalence. **(B)** Seed-matched AUROC and AUPRC on the held-out test set. **(C)** Changes in AUPRC and AUROC for each ablation relative to the corresponding same-seed MELD model. **(D)** Held-out AUPRC as a function of trainable parameter count across the architecture-capacity sweep. The dashed line connects the empirically Pareto-optimal architectures based on five-seed mean AUPRC, where parameter count is minimized and AUPRC is maximized. Transformer-only, CNN-only, and MELD lie on this frontier. Points denote individual seeds; summary error bars show mean ± sample standard deviation.

The encoder controls also highlight the importance of convolutional feature extraction for variant-effect prediction. The CNN-only model remained competitive, although adding a single Transformer layer produced small relative improvements of 0.22% in AUROC, 0.35% in AUPRC, 0.64% in F1, and 0.87% in MCC, together with relative reductions of 11.9% in Brier score and 17.9% in ECE, as shown in Supplementary Table S4. In contrast, the Transformer-only model performed substantially worse across all metrics, suggesting that local convolutional representations are imperative for this task. Based on these results, MELD (5CNN+1T) is retained as the primary architecture because it provides the strongest overall performance while remaining compact and parameter-efficient.

### 5.2 Hyperparameter sensitivity analysis

We performed a grouped hyperparameter sensitivity analysis on the compact 5CNN+1T MELD architecture, comprising 36 independently trained configurations that varied optimizer settings, delta-regularizer weights, adaptive-alpha settings, data augmentation, and dropout. Each configuration was trained and selected independently on validation AUPRC, with threshold-dependent metrics using a threshold set by maximizing validation F1 and applied once to the held-out test split; no test metric was used for configuration, checkpoint, or threshold selection. The analysis therefore represents grouped sensitivity testing rather than sequential joint optimization. Full per-configuration results are reported in Supplementary Table S5 and shown in Supplementary Fig. S1A–E.

The highest validation AUPRC was obtained with reverse-complement augmentation probability 0.5 and no positional shift, which slightly exceeded the matched standard-shift ([−8, 8]) control on validation and held-out test (Supplementary Table S6; Supplementary Fig. S1F). However, this sweep used a single training seed and is treated as a sensitivity analysis rather than a robustness estimate. The five-seed compact MELD configuration with reverse-complement probability 0.5 and positional shifts in [−8, 8] remains the primary model reported in this study.

Across the sweep, only optimizer settings materially affected performance: the three highest-learning-rate configurations collapsed to AUPRC ≈ 0.75, whereas all delta-regularizer, alpha, and dropout configurations were tightly clustered near AUPRC 0.94 (Supplementary Table S5). This indicates the model is robust to most hyperparameter choices within the tested ranges, with learning rate being the main exception.

### 5.3 Core model and ablation analysis

To test whether direct paired WT/mutant training improves over post-hoc mutation scoring, we compare MELD against a same-data, same-backbone ISM-style baseline. The ISM baseline trains an absolute single-sequence model and computes the variant score after training as

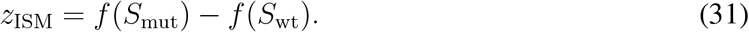

In contrast, MELD directly trains on the WT–mutant differential:

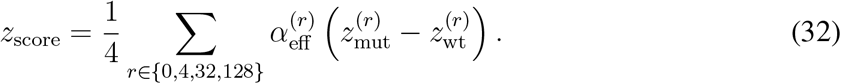

Figure 3A–C summarizes the primary MELD–ISM comparison and component-ablation results, with complete five-seed metrics reported in Supplementary Table S7. For each seed, threshold-dependent metrics were computed using a threshold selected on the validation split by maximizing validation F1 and then applied once to the held-out test split. The architecture-capacity analysis is summarized separately in Fig. 3D, with complete metrics reported in Supplementary Table S4.

MELD substantially outperforms the same-backbone post-hoc ISM baseline (AUROC 0.986 vs 0.800; AUPRC 0.934 vs 0.700). The no-mutation-boost, no-RC, no-delta-regularizer, and 256-bp context ablations all reduce performance, indicating that mutation-aware hidden-state modulation, reverse-complement augmentation and test-time averaging, delta regularization, and the full 1,025-bp variant-centered sequence context each contribute to the final model.

The no-alpha ablation remains highly competitive and slightly exceeds MELD on validation-selected-threshold F1 and MCC, whereas MELD has slightly higher AUROC (0.986 vs 0.983) and AUPRC (0.934 vs 0.931), as reported in Supplementary Table S7. Taken together, the full paired MELD configuration provides a substantial performance advantage over the tested WT-anchored post-hoc baseline, while adaptive alpha primarily affects ranking performance rather than uniformly improving all thresholded metrics. Residual-alpha variants perform similarly to MELD, suggesting that performance is relatively insensitive to the tested values of *α*_max_.

### 5.4 Performance across variant types and molecular consequences

We examined MELD’s performance across recorded variant types and transcript-aware molecular consequences in the 54,513-variant ClinVar held-out test set (Fig. 4). The recorded variant-type groups comprised 48,835 single-nucleotide variants (SNVs), 3,587 deletions, 1,993 insertions and 98 multinucleotide variants (MNVs). MELD achieved an AUPRC of 0.8916 ± 0.0045 for SNVs, 0.9748 ± 0.0057 for deletions and 0.9710 ± 0.0107 for insertions (Fig. 4B). The corresponding pathogenic fractions were 10.2%, 62.4% and 56.4%, respectively, providing context for the high precision–recall performance within these groups. AUPRC was 0.8585 ± 0.0079 for MNVs, although this estimate was based on only 98 variants.

**Fig 4:**
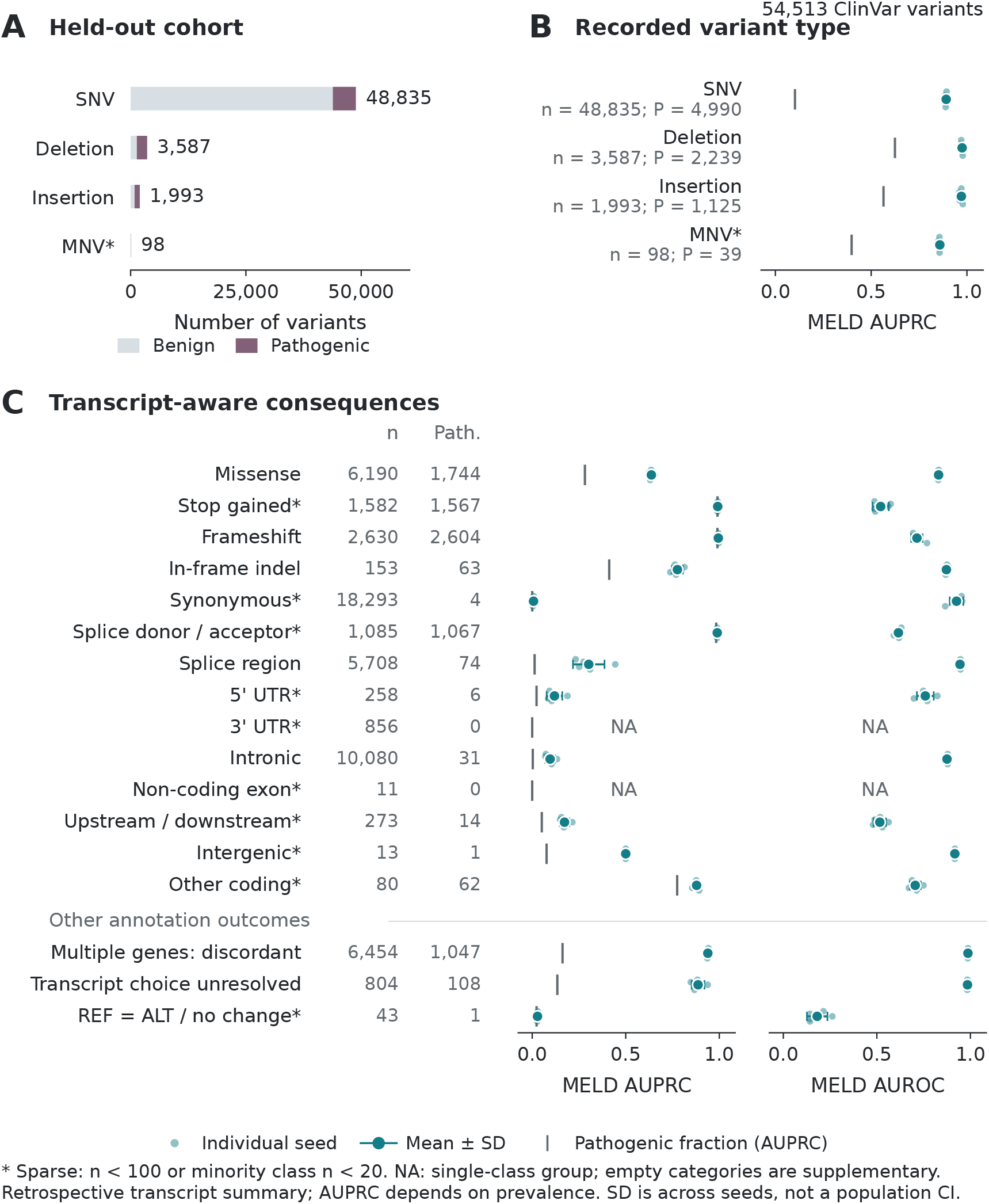
MELD performance across variant types and transcript-aware molecular consequences. **(A)** Distribution of benign and pathogenic variants across recorded variant types in the ClinVar held-out test set. **(B)** AUPRC by variant type. **(C)** AUPRC and AUROC by transcript-aware consequence category. Total and pathogenic variant counts are indicated by *n* and *P* (Path.), respectively. Small points represent individual training seeds; large markers and error bars indicate the mean ± sample standard deviation across five seeds. Grey vertical markers indicate the pathogenic fraction in each group on the AUPRC axis. “*” denote groups with fewer than 100 variants or fewer than 20 examples in either class. NA indicates a single-class group. Other annotation outcomes are shown separately; categories containing no variants are omitted.

Performance varied across molecular consequence categories (Fig. 4C). MELD distinguished pathogenic from benign variants within the missense group, which contained 6,190 variants, including 1,744 pathogenic examples, achieving AUROC 0.8300 ± 0.0034 and AUPRC 0.6371 ± 0.0115. Discrimination was also observed among in-frame indels, comprising 153 variants with 63 pathogenic examples, with AUROC 0.8717 ± 0.0081 and AUPRC 0.7762 ± 0.0296. Splice-region variants showed high AUROC (0.9442 ± 0.0037) and AUPRC above the pathogenic fraction (0.3023 ± 0.0842), despite containing only 74 pathogenic examples among 5,708 variants (1.30%). Intronic variants exhibited a similar pattern of high AUROC (0.8748 ± 0.0062) but lower absolute AUPRC (0.0957 ± 0.0254), with 31 pathogenic examples among 10,080 variants. These findings indicate within-category discrimination across several molecular consequence classes, although precision-recall performance varied considerably.

Class composition was important when interpreting the remaining groups. Pathogenic variants accounted for 99.1% of stop-gained, 99.0% of frameshift and 98.3% of splice donor/acceptor variants. Although AUPRC approached unity in these groups, AUROC indicated more limited within-category discrimination, particularly for stop-gained variants, where performance was close to chance. The small numbers of benign examples further limited interpretation of these estimates. Conversely, the synonymous and intergenic groups contained only four and one pathogenic examples, respectively, limiting interpretation of their high AUROC values. AUROC and AUPRC were not reported for the 3^′^ UTR and non-coding exon groups because these subsets contained no pathogenic variants. Overall, MELD showed discrimination across several major variant and consequence classes, while the stratified analysis revealed limitations not apparent from the aggregate performance alone.

### 5.5 Shared-subset benchmark for external foundation-model baselines

For compute-intensive external baselines, we constructed a deterministic shared subset with 200k training variants, 20k validation variants, and 20k held-out test variants. This benchmark is reported separately from the full ClinVar benchmark and is used only for fair comparison against external long-context or foundation-model baselines. The shared subset preserves the original split boundaries: training variants are sampled from the original training chromosomes only, validation variants from the original validation split, and test variants from the original held-out test split. No variants are reshuffled or moved across splits. The same fixed subset is used across all five random seeds, so the reported standard deviations reflect stochastic training variability rather than dataset resampling.

As shown in Supplementary Table S8, both MELD and its no-alpha ablation achieved higher AUROC, AUPRC, F1, and MCC when trained on the full ClinVar dataset than when trained on the shared subset, while being evaluated on the same fixed test set. These results demonstrate the benefit of additional training data for both configurations.

Across the controlled shared-subset benchmark, MELD achieved the strongest performance among the reported models across AUROC, AUPRC, F1, and MCC. MELD reached an AUROC of 0.9674 ± 0.0007 and an AUPRC of 0.8883 ± 0.0022, compared with 0.9356 ± 0.0008 and 0.8034±0.0014 for Nucleotide Transformer,^21^ the strongest frozen-representation baseline on these ranking metrics. AlphaGenome^7^ followed with an AUROC of 0.9237 ± 0.0010 and an AUPRC of 0.7922 ± 0.0033 as illustrated in Table 2 and Supplementary Fig. S2.

**Table 2:** Shared-subset ClinVar comparison on the fixed 20,000-variant held-out test set. Values are five-seed mean ± standard deviation. MELD is trained end-to-end; external backbones remain frozen to generate embeddings while the corresponding prediction heads are trained independently. Parameter counts refer to MELD’s complete model and the external backbones, excluding their prediction heads. Additional metrics are reported in Supplementary Tables S9 and S10.

| **Model** | **Parameters** | **AUROC** | **AUPRC** | **F1** | **MCC** |
|---|---|---|---|---|---|
| MELD | 1.92 M | **0.9674 ± 0.0007** | **0.8883 ± 0.0022** | **0.8030 ± 0.0046** | **0.7680 ± 0.0053** |
| Enformer<sup>19</sup> | 251.28 M | 0.7230 ± 0.0056 | 0.4196 ± 0.0027 | 0.5038 ± 0.0124 | 0.4200 ± 0.0218 |
| Borzoi<sup>20</sup> | 171.30 M | 0.8447 ± 0.0049 | 0.6200 ± 0.0053 | 0.5498 ± 0.0099 | 0.4801 ± 0.0159 |
| AlphaGenome<sup>7</sup> | 450.56 M | 0.9237 ± 0.0010 | 0.7922 ± 0.0033 | 0.7161 ± 0.0026 | 0.6676 ± 0.0037 |
| Nucleotide Transformer<sup>21</sup> | 2.55 B | 0.9356 ± 0.0008 | 0.8034 ± 0.0014 | 0.7220 ± 0.0024 | 0.6729 ± 0.0026 |
| DNABERT-2<sup>22</sup> | 117.07 M | 0.7855 ± 0.0007 | 0.5496 ± 0.0015 | 0.5087 ± 0.0019 | 0.4251 ± 0.0058 |
| Caduceus<sup>23</sup> | 7.73 M | 0.8404 ± 0.0023 | 0.5897 ± 0.0048 | 0.5300 ± 0.0053 | 0.4383 ± 0.0082 |
| HyenaDNA<sup>24</sup> | 6.55 M | 0.6746 ± 0.0044 | 0.3974 ± 0.0046 | 0.4819 ± 0.0002 | 0.4205 ± 0.0002 |

These results indicate that, on this controlled shared-subset benchmark, an architecture 3–1,300x smaller than the external baselines, trained directly on paired WT/mutant sequences can outper-form larger pretrained genomic sequence models coupled to downstream classifiers.

### 5.9 End-to-end fine-tuning of DNA language models

To complement the frozen-representation benchmark, we fully fine-tuned three relatively smaller foundation models, DNABERT-2,^22^ Caduceus^23^ and HyenaDNA,^24^ end-to-end on the full ClinVar training split and evaluated them on the same 54,513-variant held-out test set used for MELD (Table 3). Each backbone used the DeltaEmbedding head of the frozen baselines and was trained with five seeds, with checkpoints selected by validation AUPRC and classification thresholds selected by maximizing validation F1. DNABERT-2, the largest of the three (≈117M parameters), achieved a held-out AUROC of 0.9692 ± 0.0008 and AUPRC of 0.9066 ± 0.0006, compared with 0.9859 ± 0.0007 and 0.9342 ± 0.0052 for MELD. The corresponding values were 0.9561 ± 0.0018 and 0.8636 ± 0.0041 for Caduceus, and 0.9271 ± 0.0042 and 0.7397 ± 0.0129 for HyenaDNA.

**Table 3:** End-to-end fine-tuned DNA language models compared with MELD on the full ClinVar standard held-out test set (54,513 variants). All models were trained on the full ClinVar training split. Values are five-seed mean ± standard deviation; checkpoints were selected using validation AUPRC, and F1 and MCC use the threshold that maximizes validation F1. Parameter counts refer to MELD’s complete model and to the pretrained backbones, excluding their prediction heads. All metrics are reported in Supplementary Table S11.

| **Model** | **Parameters** | **AUROC** | **AUPRC** | **F1** | **MCC** |
|---|---|---|---|---|---|
| MELD | 1.92 M | **0.9859 ± 0.0007** | **0.9342 ± 0.0052** | **0.8598 ± 0.0047** | **0.8358 ± 0.0050** |
| DNABERT-2<sup>22</sup> (FT) | 117.07 M | 0.9692 ± 0.0008 | 0.9066 ± 0.0006 | 0.8293 ± 0.0024 | 0.8038 ± 0.0025 |
| Caduceus<sup>23</sup> (FT) | 7.73 M | 0.9561 ± 0.0018 | 0.8636 ± 0.0041 | 0.7817 ± 0.0035 | 0.7446 ± 0.0033 |
| HyenaDNA<sup>24</sup> (FT) | 6.55 M | 0.9271 ± 0.0042 | 0.7397 ± 0.0129 | 0.7029 ± 0.0074 | 0.6474 ± 0.0086 |

None of the fine-tuned models outperformed MELD on AUROC, AUPRC, F1 or MCC, although fine-tuned DNABERT-2 was better calibrated (Brier 0.0423 vs 0.0472; ECE 0.0295 vs 0.0711; Supplementary Table S11). Taken together, these experiments extend the external-model evaluation to end-to-end supervised adaptation rather than frozen-representation transfer alone.

### 5.7 Comparison with precomputed genome-wide scores

On the 48,795 held-out SNVs scored by all methods, CADD v1.7 achieved the highest performance on every metric (AUROC 0.9890, AUPRC 0.9400, F1 0.8780, MCC 0.8640), exceeding MELD (AUROC 0.9823 ± 0.0007, AUPRC 0.8941 ± 0.0032, F1 0.8041 ± 0.0028, MCC 0.7830 ± 0.0031; Table 4). MELD ranked second, above GPN-MSA (AUROC 0.9604, AUPRC 0.8350) and both conservation scores, phyloP (AUROC 0.9199, AUPRC 0.6773) and phastCons (AUROC 0.8537, AUPRC 0.3345). The same ordering held on the shared-subset test set, where MELD trained on full ClinVar reached an AUROC of 0.9821 ± 0.0005 against 0.9896 for CADD. MELD trained on the 200k shared subset reached an SNV AUROC of 0.9573 ± 0.0010, slightly below GPN-MSA (0.9626).

**Table 4:** Comparison with precomputed genome-wide scores on matched single-nucleotide variants. Each block is restricted to test SNVs scored by both CADD v1.7 and GPN-MSA. MELD values are five-seed mean ± standard deviation; precomputed scores are fixed lookups. F1 and MCC use the threshold that maximizes F1 on the matching validation SNVs. MELD values differ from Table 2 and Supplementary Table S3 because indels and MNVs are excluded here. Bold marks the best value in each block.

| **Model** | **AUROC** | **AUPRC** | **F1** | **MCC** |
|---|---|---|---|---|
| *Full ClinVar test set (n = 48,795 SNVs; 4,989 pathogenic)* |  |  |  |  |
| MELD | 0.9823 ± 0.0007 | 0.8941 ± 0.0032 | 0.8041 ± 0.0028 | 0.7830 ± 0.0031 |
| CADD v1.7<sup>32</sup> | **0.9890** | **0.9400** | **0.8780** | **0.8640** |
| GPN-MSA<sup>33</sup> | 0.9604 | 0.8350 | 0.7880 | 0.7635 |
| phyloP-100way<sup>34</sup> | 0.9199 | 0.6773 | 0.6649 | 0.6281 |
| phastCons-100way<sup>35</sup> | 0.8537 | 0.3345 | 0.4930 | 0.4516 |
| *Shared-subset test set (n = 17,908 SNVs; 1,830 pathogenic)* |  |  |  |  |
| MELD (full ClinVar training) | 0.9821 ± 0.0005 | 0.8921 ± 0.0034 | 0.7991 ± 0.0087 | 0.7793 ± 0.0071 |
| MELD (shared-subset training) | 0.9573 ± 0.0010 | 0.8124 ± 0.0027 | 0.7222 ± 0.0044 | 0.6951 ± 0.0041 |
| CADD v1.7<sup>32</sup> | **0.9896** | **0.9448** | **0.8818** | **0.8683** |
| GPN-MSA<sup>33</sup> | 0.9626 | 0.8317 | 0.7798 | 0.7544 |
| phyloP-100way<sup>34</sup> | 0.9249 | 0.6789 | 0.6633 | 0.6263 |
| phastCons-100way<sup>35</sup> | 0.8566 | 0.3384 | 0.4972 | 0.4561 |

Restricting to SNVs lowers MELD’s AUROC on the full test set from 0.986 to 0.982. The excluded insertions, deletions and MNVs cannot be scored by CADD or GPN-MSA from their precomputed files, and MELD efficiently classifies them with higher precision (Fig. 4B).

CADD is not trained on ClinVar labels, but it integrates more than a hundred annotations, including cross-species conservation, gene and protein-level features, and protein language model scores.^32^ Pathogenic ClinVar SNVs are concentrated at conserved coding and canonical splice positions, where these annotations are most informative. In addition, computational predictions are used as supporting evidence during clinical variant classification, which makes ClinVar labels partly circular with respect to such scores.^38^ MELD, by contrast, uses only the 1,025-bp DNA sequence around the variant, without alignments or external annotations or prior knowledge incorporation. These results show that aggregate ClinVar performance does not establish a significant advantage for MELD over integrative annotation-based scores, and that evaluation on label sources independent of clinical curation is needed to separate the two.

### 5.8 Data-efficiency and training-set scaling

We next examined whether the advantage of directly learning the paired WT/mutant differential is maintained across different amounts of supervised training data. MELD and the same-backbone ISM baseline were trained using identical nested ClinVar subsets ranging from 10,000 examples to the complete 1,482,739-variant training set, while keeping the validation and held-out test sets fixed.

As shown in Fig. 5, both models benefited from increasing amounts of training data, but MELD consistently achieved higher performance across all evaluated training-set sizes. Complete five-seed metrics are reported in Supplementary Table S12. With 10,000 training variants, MELD achieved an AUROC of 0.8090 ± 0.0031 and an AUPRC of 0.5448 ± 0.0068, compared with 0.5092 ± 0.0103 and 0.2388 ± 0.0112, respectively, for ISM.

**Fig 5:**
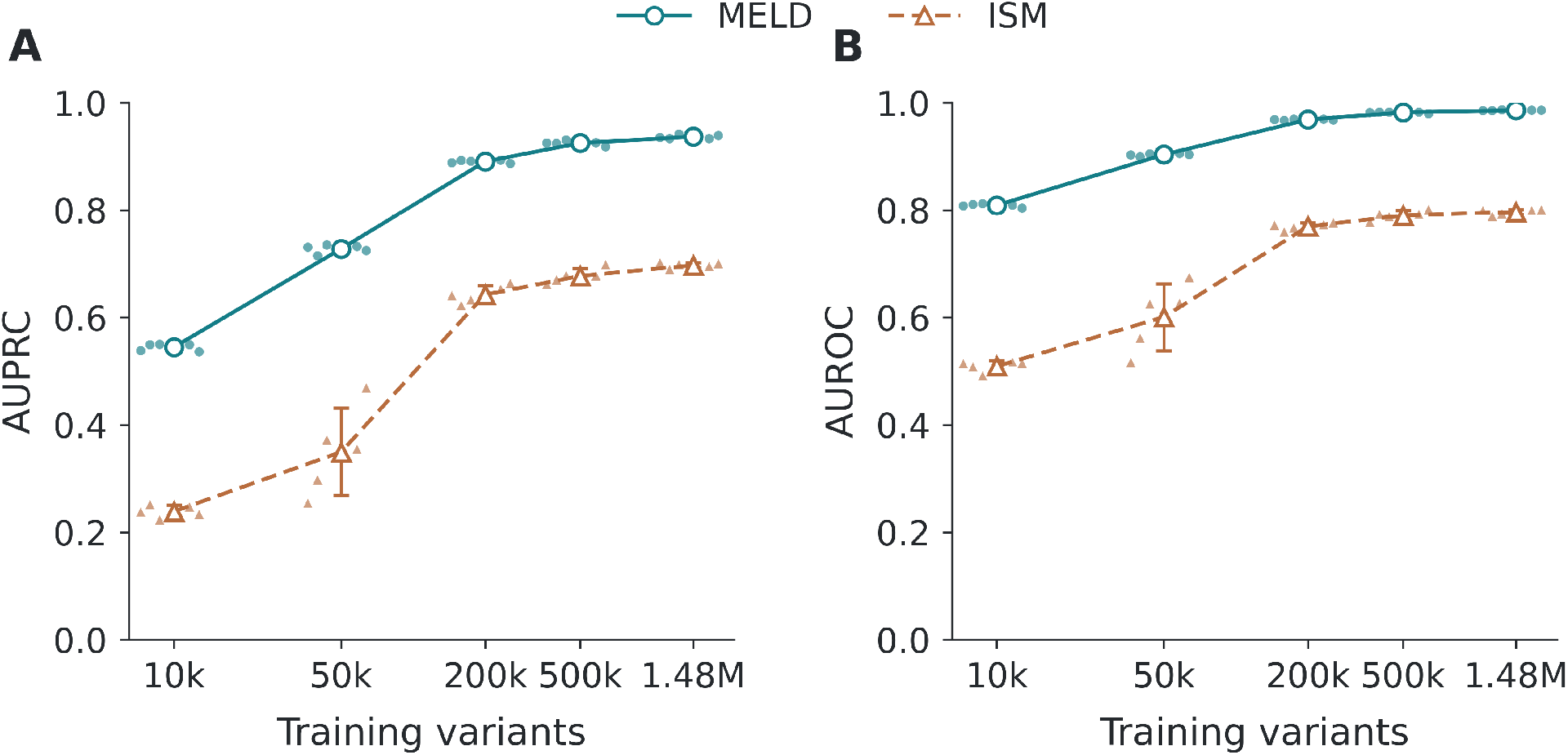
Data-efficiency of MELD and the same-backbone ISM baseline. **(A)** AUPRC and **(B)** AUROC on the fixed ClinVar held-out test set across increasing training-set sizes. Small points denote individual seeds, horizontally offset for visibility; larger markers and error bars show the five-seed mean ± sample standard deviation.

ISM performance improved substantially as the training set increased, with the largest gains occurring between 50,000 and 200,000 variants. Using the complete training set, ISM reached an AUROC of 0.7963 ± 0.0048, an AUPRC of 0.6973 ± 0.0049, an F1 score of 0.6818 ± 0.0058, and an MCC of 0.6537 ± 0.0076. MELD maintained a clear advantage at full scale, achieving an AUROC of 0.9861 ± 0.0004, an AUPRC of 0.9367 ± 0.0037, an F1 score of 0.8619 ± 0.0034, and an MCC of 0.8379 ± 0.0036.

Notably, MELD trained on only 50,000 variants already exceeded full-data ISM in both AU-ROC (0.9034 versus 0.7963) and AUPRC (0.7279 versus 0.6973). These results indicate that explicitly learning the paired WT/mutant differential provides a considerable advantage in data efficiency, with MELD maintaining higher predictive performance across the complete range of training-set sizes.

### 5.9 Quantitative BRCA1 mutation-effect regression

We next evaluated whether MELD can extend from binary pathogenicity prediction to continuous experimental mutation-effect prediction using the BRCA1 saturation genome editing benchmark. MELD-DeltaEmbedding was compared with the same-backbone ISM regression baseline and frozen pretrained genomic encoders using the common DeltaEmbedding formulation. External model configurations were selected using validation Spearman correlation and fixed held-out evaluation.

Training MELD directly on the BRCA1 regression benchmark from random initialization already outperformed the same-backbone ISM baseline in both Pearson *r* and Spearman *ρ*, reaching 0.2530 ± 0.0182 and 0.3175 ± 0.0225, respectively. Freezing the strict ClinVar-pretrained MELD representation and training only the BRCA1 regression head substantially improved performance, reaching a Pearson *r* of 0.4720 ± 0.0132 and a Spearman *ρ* of 0.4172 ± 0.0117. End-to-end fine-tuning from the same strict ClinVar-pretrained initialization provided an additional improvement, increasing Pearson *r* to 0.5130 ± 0.0096 and Spearman *ρ* to 0.4501 ± 0.0133. Strict transfer also reduced RMSE to 0.6982 ± 0.0075 and MAE to 0.5576 ± 0.0126, as observed in Table 5 and Supplementary Fig. S3A-C.

**Table 5:**
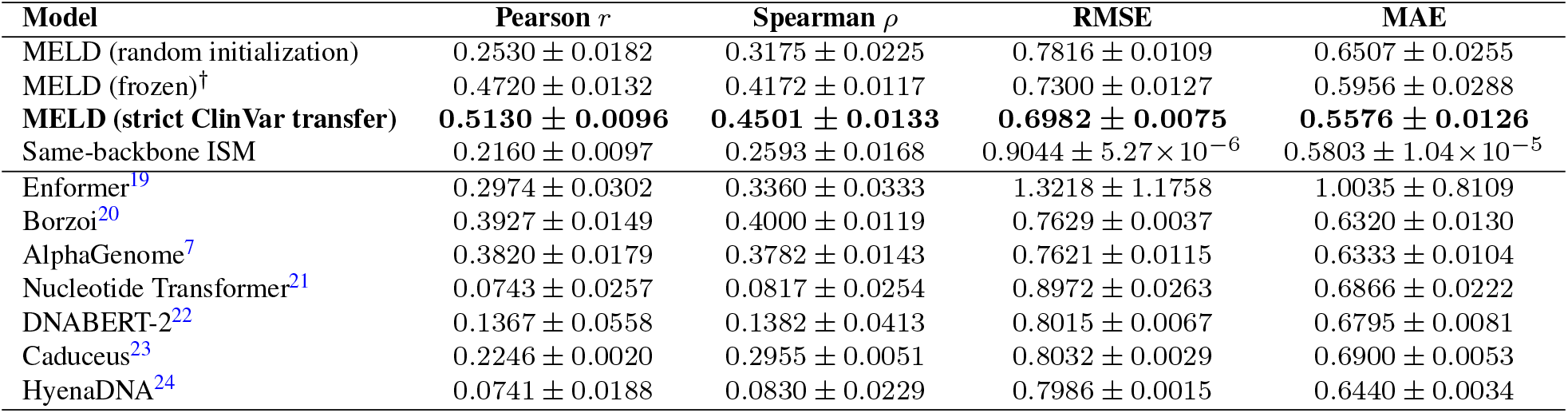
Quantitative BRCA1 mutation-effect prediction on the fixed 466-variant held-out test set. Values are reported as five-seed mean ± sample standard deviation. MELD and the frozen external pretrained encoders use the DeltaEmbedding regression formulation, whereas the same-backbone ISM baseline uses post-hoc mutant-minus-wild-type scoring. ^†^The frozen MELD condition uses the strict ClinVar-pretrained representation with the encoder frozen and only the BRCA1 DeltaEmbedding regression head trained; this condition was added post hoc as a protocol-matched frozen-representation control.

| **Model** | **Pearson *r*** | **Spearman *ρ*** | **RMSE** | **MAE** |
|---|---|---|---|---|
| MELD (random initialization) | 0.2530 ± 0.0182 | 0.3175 ± 0.0225 | 0.7816 ± 0.0109 | 0.6507 ± 0.0255 |
| MELD (frozen)<sup>†</sup> | 0.4720 ± 0.0132 | 0.4172 ± 0.0117 | 0.7300 ± 0.0127 | 0.5956 ± 0.0288 |
| **MELD (strict ClinVar transfer)** | **0.5130 ± 0.0096** | **0.4501 ± 0.0133** | **0.6982 ± 0.0075** | **0.5576 ± 0.0126** |
| Same-backbone ISM | 0.2160 ± 0.0097 | 0.2593 ± 0.0168 | 0.9044 ± 5.27 × 10<sup>−6</sup> | 0.5803 ± 1.04 × 10<sup>−5</sup> |
| Enformer<sup>19</sup> | 0.2974 ± 0.0302 | 0.3360 ± 0.0333 | 1.3218 ± 1.1758 | 1.0035 ± 0.8109 |
| Borzoi<sup>20</sup> | 0.3927 ± 0.0149 | 0.4000 ± 0.0119 | 0.7629 ± 0.0037 | 0.6320 ± 0.0130 |
| AlphaGenome<sup>7</sup> | 0.3820 ± 0.0179 | 0.3782 ± 0.0143 | 0.7621 ± 0.0115 | 0.6333 ± 0.0104 |
| Nucleotide Transformer<sup>21</sup> | 0.0743 ± 0.0257 | 0.0817 ± 0.0254 | 0.8972 ± 0.0263 | 0.6866 ± 0.0222 |
| DNABERT-2<sup>22</sup> | 0.1367 ± 0.0558 | 0.1382 ± 0.0413 | 0.8015 ± 0.0067 | 0.6795 ± 0.0081 |
| Caduceus<sup>23</sup> | 0.2246 ± 0.0020 | 0.2955 ± 0.0051 | 0.8032 ± 0.0029 | 0.6900 ± 0.0053 |
| HyenaDNA<sup>24</sup> | 0.0741 ± 0.0188 | 0.0830 ± 0.0229 | 0.7986 ± 0.0015 | 0.6440 ± 0.0034 |

Under a protocol matched to the external pretrained baselines, frozen MELD achieved stronger mean held-out performance than all evaluated frozen genomic encoders across the four reported regression metrics. Borzoi^20^ was the strongest external baseline in rank correlation, reaching a Spearman *ρ* of 0.4000 ± 0.0119, compared with 0.4172 ± 0.0117 for frozen MELD. Frozen MELD also achieved a Pearson *r* of 0.4720 ± 0.0132, compared with 0.3927 ± 0.0149 for Borzoi^20^ and 0.3820 ± 0.0179 for AlphaGenome,^7^ while obtaining lower RMSE and MAE than every external baseline. End-to-end strict ClinVar transfer further improved all four metrics and produced the strongest overall performance. Together, these results show that mutation-specific ClinVar supervision produces transferable MELD representations even when the pretrained encoder is held fixed, while BRCA1-specific end-to-end fine-tuning provides additional benefit.

A sensitivity analysis using progressively stricter exclusion of BRCA1-related ClinVar training examples is reported in Supplementary Table S13 and Supplementary Fig. S3D. Although excluding only exact BRCA1 target alleles yielded stronger validation performance, this setting still permitted ClinVar examples with overlapping local sequence contexts. We therefore retained the stricter window-excluded configuration for confirmatory evaluation because it removes both exact target-allele overlap and overlapping input-window contexts.

### 5.10 Structural context of BRCA1 variant-effect predictions

To relate the quantitative predictions to protein structure, we mapped held-out BRCA1 variants onto annotated functional regions and the experimental tandem BRCT structure (PDB: 1JNX;^39^). The held-out coverage was confined to the BRCT regions (Fig. 6A). For the strict ClinVar-transfer model illustrated using seed 42, predicted and observed SGE damage scores showed a positive association across 466 variants, with Pearson *r* = 0.518 and Spearman *ρ* = 0.452 (Fig. 6B).

**Fig 6:**
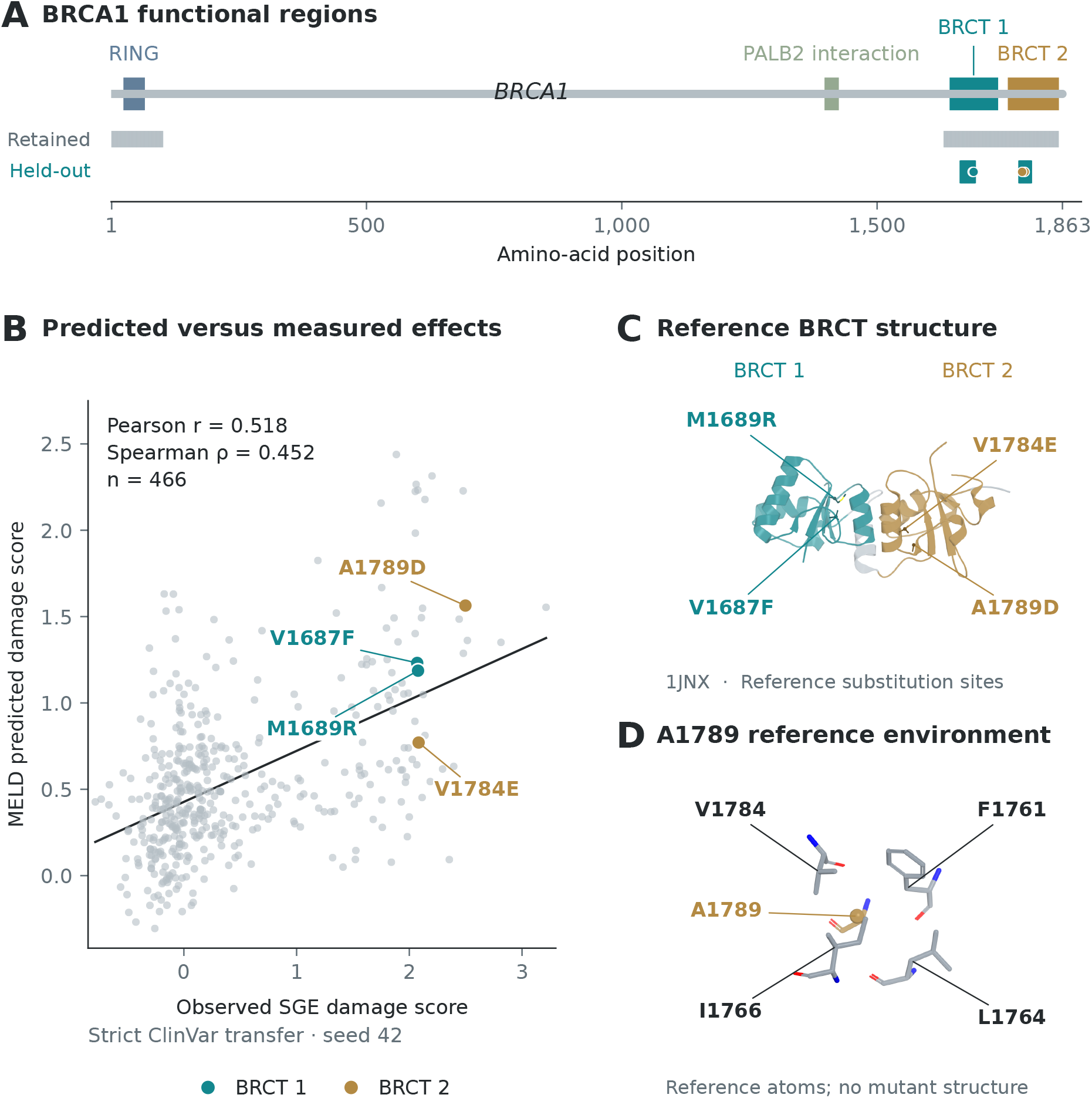
Structural context of BRCA1 variant-effect predictions. **(A)** BRCA1 domain organization and protein regions represented by the retained and held-out benchmark variants. **(B)** Predicted versus observed SGE damage scores for 466 held-out variants using the strict ClinVar-transfer MELD model with seed 42. Higher scores indicate greater functional impairment. The solid line shows a descriptive linear fit; selected substitutions are highlighted by BRCT domain. **(C)** Sites of V1687F, M1689R, V1784E and A1789D mapped onto the experimental reference BRCT structure (PDB: 1JNX). **(D)** Local reference-residue environment around A1789, showing selected neighbouring residues. Structural views depict reference coordinates, not mutant conformations.

Four illustrative missense substitutions were examined in structural context: V1687F and M1689R in BRCT1, and V1784E and A1789D in BRCT2 (Fig. 6C). All four had elevated measured SGE damage scores and positive predicted damage scores, although MELD underestimated their measured effect magnitudes. These examples illustrate the correspondence between predicted effects and experimentally measured impairment, while also showing the remaining limitations of quantitative prediction.

Inspection of the reference structure placed A1789 in a hydrophobic environment containing F1761, L1764, I1766 and V1784 (Fig. 6D). Replacing alanine with an acidic aspartate could perturb local packing, providing a plausible structural interpretation of A1789D. Consistent with the importance of this position, a recent study identified A1789 as a BRCT hydrophobic-core residue and demonstrated impaired partner interactions and homologous recombination for A1789L.^40^

Those findings concern a different substitution and support the functional importance of the residue rather than establishing the mechanism of A1789D. Together, the structural mapping and SGE comparison connect MELD predictions to functional variation within the BRCT region, without attributing the observed effects to a directly demonstrated conformational change.

## 6 Discussion & Conclusion

This work tests a simple premise: that a variant’s effect is better learned directly than recovered after the fact. Our proposed approach, MELD, places the wild-type (WT) versus mutant logit differential Δ*z* inside the supervised objective, and under matched data, split, and backbone this change alone raised ClinVar AUROC from 0.800 (post-hoc ISM) to 0.986, and AUPRC from 0.700 to 0.934. Because the ISM baseline shares MELD’s encoder family and training data, this gap is attributable to the training formulation rather than to model capacity or data. The advantage was most pronounced in low-data regimes: with 10,000 training variants MELD reached an AUROC of 0.809 against 0.509 for ISM, and with 50,000 variants it already exceeded the full-data ISM baseline (AUROC 0.903 vs. 0.796). These findings indicate that the difference signal is a more efficient learning target than absolute single-sequence labels, from which sensitivity to nucleotide-level perturbation must be acquired indirectly.

The results also show that this formulation does not require a large backbone. With approximately 1.9M parameters and a 1,025-bp window, MELD outperformed frozen Enformer,^19^ Borzoi,^20^ AlphaGenome,^7^ and four DNA language models on a controlled shared-subset benchmark (AUROC 0.967 vs. 0.936 for the strongest baseline, Nucleotide Transformer^21^), despite these models having 3-1,300× more parameters. This should be read as evidence for the value of mutation-specific supervision rather than as a claim of general superiority: the external models were trained for absolute signal prediction or self-supervised modeling, and were evaluated through a lightweight head over frozen representations. Notably, fully fine-tuned DNABERT-2^22^ (117M parameters), trained on the full ClinVar split, reached an AUROC of 0.969 against 0.986 for MELD on the same test set (Table 3), suggesting that supervised adaptation of a pretrained backbone can recover much of the same benefit, at far greater computational cost (≈ 60x).

The ablation and sensitivity analyses clarify which components matter. The mutation-mask boost, delta regularization, reverse-complement augmentation, and the full 1,025-bp context each improved performance, and convolutional feature extraction was essential, as the Transformer-only encoder performed markedly worse. Increasing encoder depth did not help and increased seed-to-seed variability, supporting the compact design. In contrast, adaptive amplification contributed modestly: the no-*α* ablation was nearly as strong (AUROC 0.983 vs. 0.986) and slightly exceeded MELD on thresholded F1 and MCC, indicating that *α* chiefly benefits ranking. Across 36 hyper-parameter configurations only very high learning rates degraded performance, indicating that the approach is not sensitive to fine-grained tuning.

The BRCA1 saturation genome editing experiments indicate that the differential representation extends beyond binary pathogenicity. Under a protocol matched to the external baselines, frozen ClinVar-pretrained MELD achieved a Spearman correlation of 0.42, exceeding Borzoi^20^ (0.40) and AlphaGenome^7^ (0.38), and end-to-end transfer raised this to 0.45. The gap between random initialization (*ρ* = 0.32) and ClinVar transfer shows that most of the quantitative gain derives from pretraining on large-scale pathogenicity labels, which supports the transferability of the learned representation. Absolute correlations remain moderate, however, and the BRCA1 test set is small.

The pattern of results across models is consistent with the biology of the benchmark. Pathogenic SNVs in ClinVar are concentrated in protein-coding sequence and at canonical splice sites, where a single substitution can introduce a premature stop codon, alter a constrained residue, or abolish exon recognition.^29, 41, 42^ These effects are largely determined by sequence close to the variant, which a compact model with a 1-kb window and convolutional feature extraction is well placed to capture. The ranking of the frozen sequence-to-function baselines may partly reflect the same mechanism. Enformer predicts regulatory and expression tracks at 128-bp resolution,^19^ Borzoi predicts RNA-seq coverage that partly reflects splicing,^20^ and AlphaGenome^7^ adds base-resolution splice-site and junction outputs,^7^ and AUROC increased in this order (0.723, 0.845, and 0.924), although representation type also differs across these models. Nucleotide Transformer, the strongest frozen baseline, was pretrained on genomes from hundreds of species,^21^ so its representations may encode evolutionary constraint, which separates pathogenic from benign variants across the genome.^34^ The precomputed scores support this reading: phyloP conservation alone reached an AUROC of 0.920 on held-out ClinVar SNVs, and CADD, which combines conservation with gene- and protein-level annotations, reached 0.989 (Table 4). Conversely, the strengths of the large sequence-to-function models lie in regulatory and long-range effects that ClinVar labels represent poorly, which is consistent with the limitations discussed below.

The BRCA1 results can be read in the same way. The saturation genome editing assay measured SNVs across the 13 exons encoding the RING and BRCT domains in HAP1 cells, where BRCA1 is essential, so the damage score is a direct readout of loss of function.^30^ In that assay, nonsense and canonical splice-site variants were almost uniformly deleterious, damaging missense variants clustered within the RING and BRCT domains, and a subset of exonic variants acted through disrupted splicing.^30^ Because ClinVar pathogenic labels largely encode the same loss-of-function mechanisms, a ClinVar-pretrained representation can transfer to BRCA1 even after overlapping training windows are removed, and the relatively strong performance of Borzoi^20^ and AlphaGenome^7^ is consistent with the contribution of splicing. Nucleotide Transformer,^21^ by contrast, ranked highly on ClinVar but correlated weakly with BRCA1 damage (*ρ* = 0.08). One possible explanation is that a representation of genome-wide constraint discriminates less well within a single gene, where most assayed positions lie in conserved coding exons. The moderate absolute correlations of all models are also expected for DNA-only inputs: the effect of a missense substitution depends on protein structure and residue-level constraint, which protein-level predictors model explicitly,^43, 44^ so combining the paired differential objective with protein-level features is a natural future extension.

Several limitations are readily addressable in future work. First, quantitative evaluation is limited to one gene and 466 test variants; additional saturation mutagenesis and MPRA datasets would establish generality and tighten confidence intervals. Second, MELD scores single variants within a 1-kb window, so long-range regulatory effects, splicing, and multi-track outputs, which are the strengths of the foundation models, are not captured; longer windows or a hybrid with a pretrained backbone are natural extensions. Third, on the SNVs it can score, the integrative CADD v1.7 score exceeded MELD on ClinVar (AUROC 0.989 vs. 0.982; Table 4), so MELD’s advantage is established against sequence-only models rather than against annotation-based genome-wide scores. Finally, ClinVar labels are subject to ascertainment bias and label noise, and a few duplicated sequence windows were retained; evaluation on independent sets such as population-frequency-derived benign variants and deep mutational scanning data would strengthen the evidence.

## Supplementary Information

### Supplementary Note 1: ClinVar benchmark audit and training subsets

The ClinVar standard benchmark contains 1,572,808 paired variant rows with a fixed, chromosome-disjoint train/validation/test split. In addition to the exact variant-, WT/mutant-pair- and sequence-window-level overlap checks described in the main text, each validation and test sequence was screened against the training set with a MinHash/LSH near-duplicate search (Supplementary Table S1). The data-efficiency experiment used deterministic nested subsets of the training split, with the validation and held-out test sets unchanged (Supplementary Table S2).

**Table S1:** Approximate sequence-similarity leakage for the ClinVar standard 1 kb benchmark. For each validation or test sequence, the audit searched for the nearest approximate match in the training set using a MinHash/LSH near-duplicate screen.

| **Mode** | **Split** | **N** | **Max sim.** | **Mean** | **Median** | **Above 90%** | **Above 95% / 99%** |
|---|---|---|---|---|---|---|---|
| WT | Test | 54,513 | 0.000 | 0.000 | 0.000 | 0 | 0 / 0 |
| WT | Validation | 35,556 | 0.000 | 0.000 | 0.000 | 0 | 0 / 0 |
| Mutant | Test | 54,513 | 0.000 | 0.000 | 0.000 | 0 | 0 / 0 |
| Mutant | Validation | 35,556 | 0.875 | ∼ 0.000 | 0.000 | 0 | 0 / 0 |
| WT/mut pair | Test | 54,513 | 0.000 | 0.000 | 0.000 | 0 | 0 / 0 |
| WT/mut pair | Validation | 35,556 | 0.000 | 0.000 | 0.000 | 0 | 0 / 0 |

**Table S2:** Nested ClinVar training subsets used for the data-efficiency experiment. The validation and held-out test sets are identical across all training sizes.

| **Training size** | **Benign** | **Pathogenic** | **Total** |
|---|---|---|---|
| 10k | 8,081 | 1,919 | 10,000 |
| 50k | 40,407 | 9,593 | 50,000 |
| 200k | 161,625 | 38,375 | 200,000 |
| 500k | 404,065 | 95,935 | 500,000 |
| Full | 1,198,246 | 284,493 | 1,482,739 |

### Supplementary Note 2: Reference model and architecture capacity

Supplementary Table S3 lists the complete held-out metrics of the reference 5CNN+1T MELD model. Supplementary Table S4 compares it with deeper encoders and with CNN-only and Transformer-only controls, all trained with the same paired objective, ClinVar split, input representation and training protocol. Held-out AUPRC against parameter count for these models is shown in Fig. 3D.

**Table S3:** Held-out test performance of the MELD reference model on the ClinVar standard benchmark. Values are reported as five-seed mean ± standard deviation.

| **Metric** | **Value** |
|---|---|
| AUROC | 0.9859 ± 0.0007 |
| AUPRC | 0.9342 ± 0.0052 |
| F1 | 0.8598 ± 0.0047 |
| MCC | 0.8358 ± 0.0050 |
| Balanced accuracy | 0.9078 ± 0.0063 |
| Accuracy | 0.9581 ± 0.0011 |
| Precision | 0.8864 ± 0.0102 |
| Recall | 0.8350 ± 0.0144 |
| Specificity | 0.9805 ± 0.0023 |
| False-positive rate | 0.0195 ± 0.0023 |
| Brier score | 0.0472 ± 0.0009 |
| Expected calibration error | 0.0711 ± 0.0014 |
| Parameters | 1,917,059 |
| Peak GPU memory | 10.22 GB |

**Table S4:** Architecture-capacity results on the ClinVar standard held-out test set using 1,025-bp variant-centered sequence windows. Values are reported as five-seed mean ± sample standard deviation.

| **Metric** | **MELD** | **5CNN+2T** | **7CNN+2T** | **7CNN+4T** | **CNN-only** | **Transformer-only** |
|---|---|---|---|---|---|---|
| CNN blocks | 5 | 5 | 7 | 7 | 5 | 0 |
| Transformer layers | 1 | 2 | 2 | 4 | 0 | 2 |
| Parameters | 1.92M | 2.12M | 2.74M | 3.14M | 1.72M | 0.56M |
| AUROC | 0.9859 ± 0.0007 | 0.9858 ± 0.0011 | 0.9799 ± 0.0031 | 0.9813 ± 0.0078 | 0.9837 ± 0.0007 | 0.8077 ± 0.0428 |
| AUPRC | 0.9342 ± 0.0052 | 0.9341 ± 0.0066 | 0.9103 ± 0.0127 | 0.9231 ± 0.0207 | 0.9309 ± 0.0024 | 0.5648 ± 0.0359 |
| F1 | 0.8598 ± 0.0047 | 0.8572 ± 0.0086 | 0.8282 ± 0.0104 | 0.8434 ± 0.0253 | 0.8543 ± 0.0015 | 0.5363 ± 0.0261 |
| MCC | 0.8358 ± 0.0050 | 0.8327 ± 0.0098 | 0.7981 ± 0.0131 | 0.8159 ± 0.0291 | 0.8286 ± 0.0024 | 0.4663 ± 0.0210 |
| Brier score | 0.0472 ± 0.0009 | 0.0496 ± 0.0032 | 0.0695 ± 0.0227 | 0.0680 ± 0.0336 | 0.0536 ± 0.0040 | 0.1998 ± 0.0417 |
| ECE | 0.0711 ± 0.0014 | 0.0769 ± 0.0064 | 0.0981 ± 0.0326 | 0.0955 ± 0.0412 | 0.0866 ± 0.0084 | 0.3098 ± 0.0584 |

### Supplementary Note 3: Hyperparameter sensitivity

The sweep comprised 36 configurations of the compact 5CNN+1T model, each trained with seed 42. Each group (augmentation, dropout, delta regularization, adaptive-alpha formulation, optimizer) was varied independently from the same base configuration; later groups did not inherit the best settings of earlier groups. Supplementary Table S5 reports held-out test metrics for every configuration, and Supplementary Fig. S1A–E plots held-out AUPRC by group. The highest validation AUPRC was obtained with reverse-complement probability 0.5 and no positional shift; Supplementary Table S6 and Supplementary Fig. S1F compare this configuration with its matched [−8, 8]-shift control. Because the sweep used a single seed, the five-seed [−8, 8] configuration is retained as the primary model.

**Fig S1:**
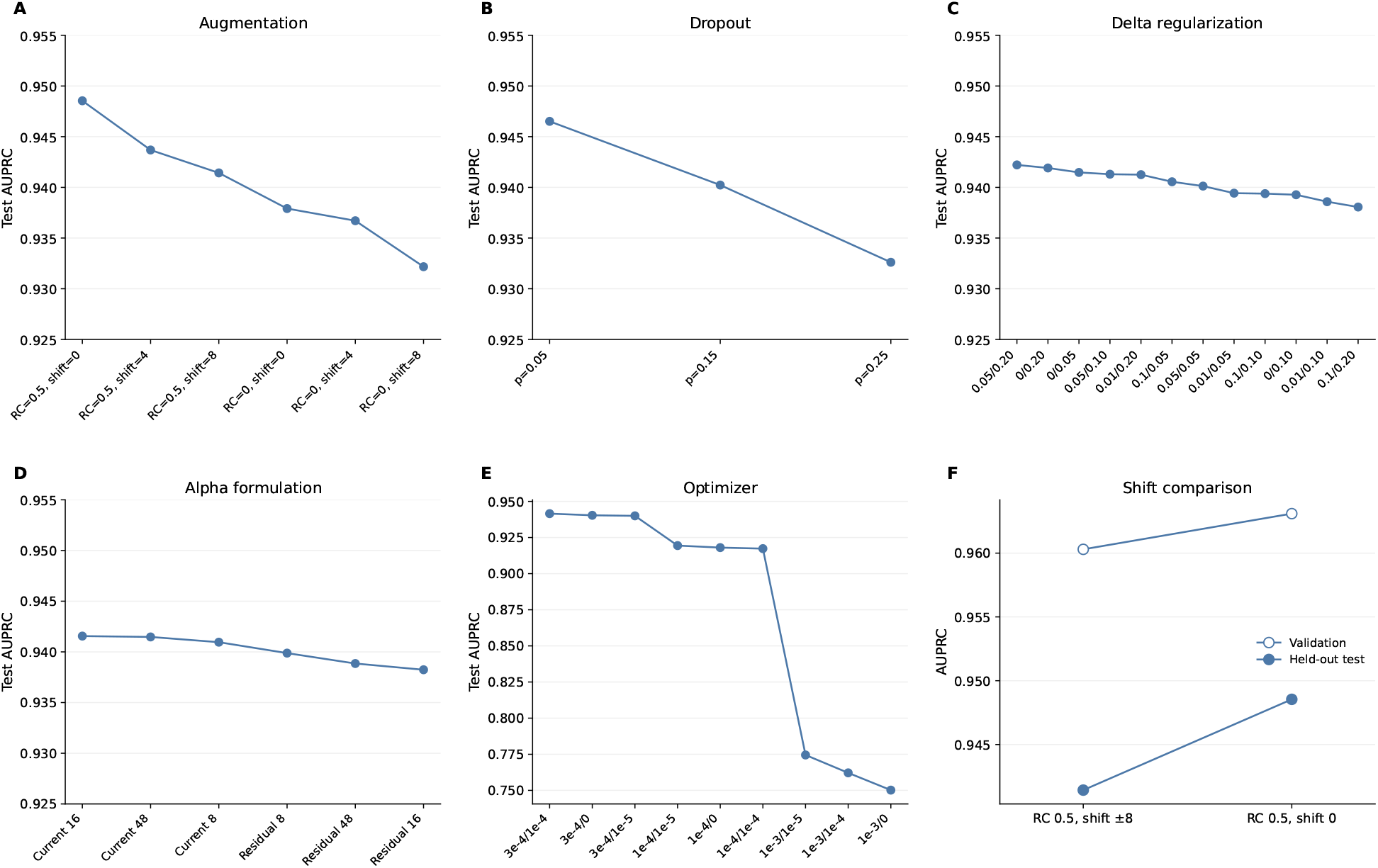
Hyperparameter sensitivity of the compact 5CNN+1T MELD model. Each point is one configuration from the 36-configuration sweep (seed 42), with the checkpoint selected on validation AUPRC. Panels **(A)**–**(E)** show held-out test AUPRC within each group, with configurations ordered from highest to lowest: **(A)** reverse-complement (RC) augmentation probability and maximum positional shift; **(B)** dropout probability; **(C)** delta-regularizer weights, labelled *λ*_Δ_*/λ*_neutral_; **(D)** adaptive-alpha formulation (current or residual) and upper bound *α*_max_; **(E)** optimizer learning rate and weight decay, labelled LR/WD. Held-out AUPRC remains between 0.93 and 0.95 across the augmentation, dropout, delta-regularization and alpha groups, whereas the three LR = 10^−3^ configurations fall to 0.75–0.77. **(F)** Validation and held-out test AUPRC for the validation-selected configuration (RC *p* = 0.5, shift 0) and its matched shift [−8, 8] control. Values for (A)–(E) are listed in Supplementary Table S5 and for (F) in Supplementary Table S6.

**Table S5:**
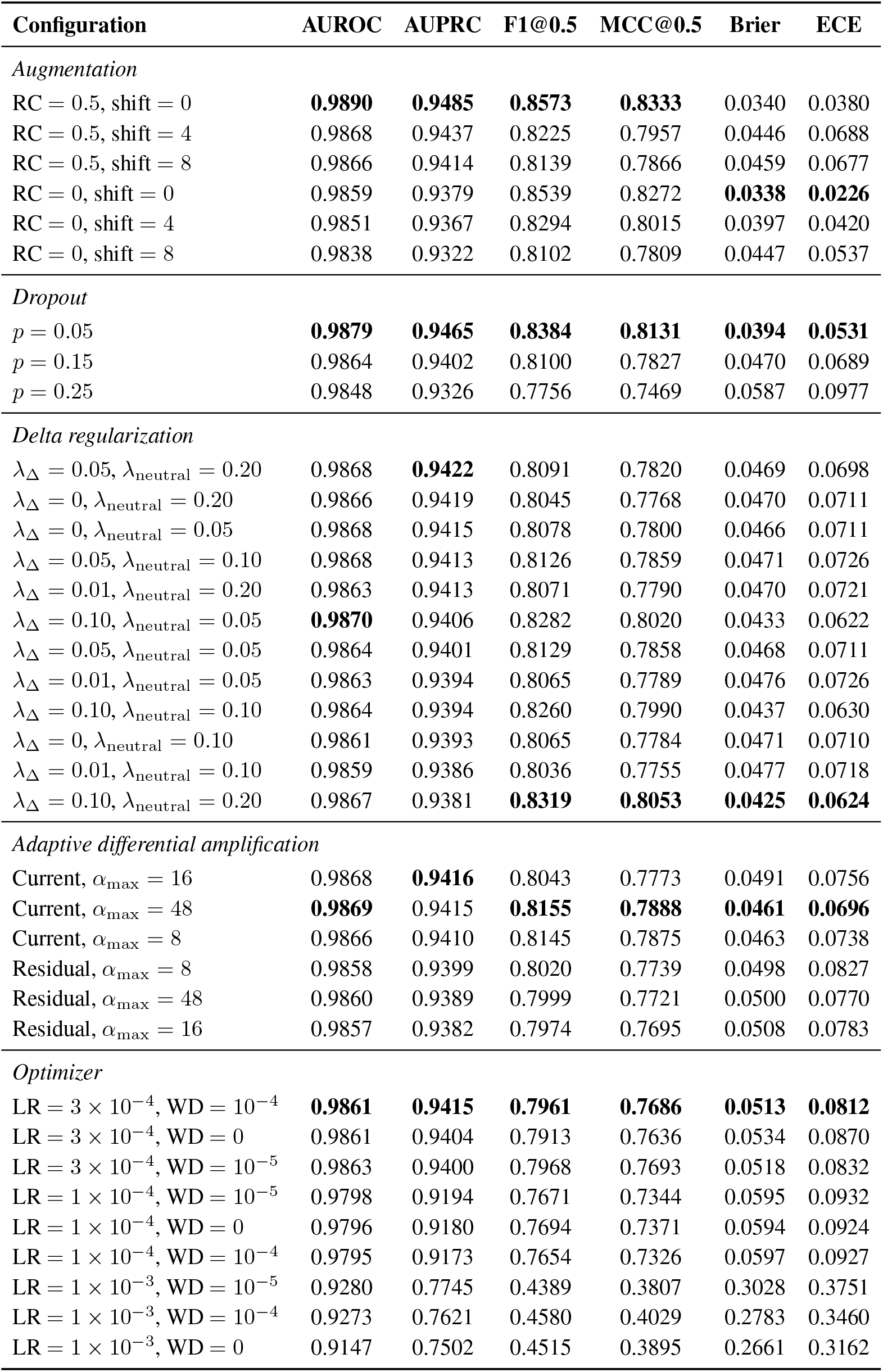
Held-out test performance across 36 hyperparameter configurations of the compact 5CNN+1T MELD model, each trained with seed 42. Checkpoints were selected using validation AUPRC. F1 and MCC are evaluated at a fixed threshold of 0.5; this differs from the validation-F1-selected thresholds used in Supplementary Table S6. Bold marks the best value within each group.

| Configuration | AUROC | AUPRC | F1@0.5 | MCC@0.5 | Brier | ECE |
|---|---|---|---|---|---|---|
| *Augmentation* |  |  |  |  |  |  |
| RC = 0.5, shift = 0 | **0.9890** | **0.9485** | **0.8573** | **0.8333** | 0.0340 | 0.0380 |
| RC = 0.5, shift = 4 | 0.9868 | 0.9437 | 0.8225 | 0.7957 | 0.0446 | 0.0688 |
| RC = 0.5, shift = 8 | 0.9866 | 0.9414 | 0.8139 | 0.7866 | 0.0459 | 0.0677 |
| RC = 0, shift = 0 | 0.9859 | 0.9379 | 0.8539 | 0.8272 | **0.0338** | **0.0226** |
| RC = 0, shift = 4 | 0.9851 | 0.9367 | 0.8294 | 0.8015 | 0.0397 | 0.0420 |
| RC = 0, shift = 8 | 0.9838 | 0.9322 | 0.8102 | 0.7809 | 0.0447 | 0.0537 |
| *Dropout* |  |  |  |  |  |  |
| *p* = 0.05 | **0.9879** | **0.9465** | **0.8384** | **0.8131** | **0.0394** | **0.0531** |
| *p* = 0.15 | 0.9864 | 0.9402 | 0.8100 | 0.7827 | 0.0470 | 0.0689 |
| *p* = 0.25 | 0.9848 | 0.9326 | 0.7756 | 0.7469 | 0.0587 | 0.0977 |
| *Delta regularization* |  |  |  |  |  |  |
| λ<sub>Δ</sub> = 0.05, λ<sub>neutral</sub> = 0.20 | 0.9868 | **0.9422** | 0.8091 | 0.7820 | 0.0469 | 0.0698 |
| λ<sub>Δ</sub> = 0, λ<sub>neutral</sub> = 0.20 | 0.9866 | 0.9419 | 0.8045 | 0.7768 | 0.0470 | 0.0711 |
| λ<sub>Δ</sub> = 0, λ<sub>neutral</sub> = 0.05 | 0.9868 | 0.9415 | 0.8078 | 0.7800 | 0.0466 | 0.0711 |
| λ<sub>Δ</sub> = 0.05, λ<sub>neutral</sub> = 0.10 | 0.9868 | 0.9413 | 0.8126 | 0.7859 | 0.0471 | 0.0726 |
| λ<sub>Δ</sub> = 0.01, λ<sub>neutral</sub> = 0.20 | 0.9863 | 0.9413 | 0.8071 | 0.7790 | 0.0470 | 0.0721 |
| λ<sub>Δ</sub> = 0.10, λ<sub>neutral</sub> = 0.05 | **0.9870** | 0.9406 | 0.8282 | 0.8020 | 0.0433 | 0.0622 |
| λ<sub>Δ</sub> = 0.05, λ<sub>neutral</sub> = 0.05 | 0.9864 | 0.9401 | 0.8129 | 0.7858 | 0.0468 | 0.0711 |
| λ<sub>Δ</sub> = 0.01, λ<sub>neutral</sub> = 0.05 | 0.9863 | 0.9394 | 0.8065 | 0.7789 | 0.0476 | 0.0726 |
| λ<sub>Δ</sub> = 0.10, λ<sub>neutral</sub> = 0.10 | 0.9864 | 0.9394 | 0.8260 | 0.7990 | 0.0437 | 0.0630 |
| λ<sub>Δ</sub> = 0, λ<sub>neutral</sub> = 0.10 | 0.9861 | 0.9393 | 0.8065 | 0.7784 | 0.0471 | 0.0710 |
| λ<sub>Δ</sub> = 0.01, λ<sub>neutral</sub> = 0.10 | 0.9859 | 0.9386 | 0.8036 | 0.7755 | 0.0477 | 0.0718 |
| λ<sub>Δ</sub> = 0.10, λ<sub>neutral</sub> = 0.20 | 0.9867 | 0.9381 | **0.8319** | **0.8053** | **0.0425** | **0.0624** |
| *Adaptive differential amplification* |  |  |  |  |  |  |
| Current, α<sub>max</sub> = 16 | 0.9868 | **0.9416** | 0.8043 | 0.7773 | 0.0491 | 0.0756 |
| Current, α<sub>max</sub> = 48 | **0.9869** | 0.9415 | **0.8155** | **0.7888** | **0.0461** | **0.0696** |
| Current, α<sub>max</sub> = 8 | 0.9866 | 0.9410 | 0.8145 | 0.7875 | 0.0463 | 0.0738 |
| Residual, α<sub>max</sub> = 8 | 0.9858 | 0.9399 | 0.8020 | 0.7739 | 0.0498 | 0.0827 |
| Residual, α<sub>max</sub> = 48 | 0.9860 | 0.9389 | 0.7999 | 0.7721 | 0.0500 | 0.0770 |
| Residual, α<sub>max</sub> = 16 | 0.9857 | 0.9382 | 0.7974 | 0.7695 | 0.0508 | 0.0783 |
| *Optimizer* |  |  |  |  |  |  |
| LR = 3 × 10<sup>−4</sup>, WD = 10<sup>−4</sup> | **0.9861** | **0.9415** | **0.7961** | **0.7686** | **0.0513** | **0.0812** |
| LR = 3 × 10<sup>−4</sup>, WD = 0 | 0.9861 | 0.9404 | 0.7913 | 0.7636 | 0.0534 | 0.0870 |
| LR = 3 × 10<sup>−4</sup>, WD = 10<sup>−5</sup> | 0.9863 | 0.9400 | 0.7968 | 0.7693 | 0.0518 | 0.0832 |
| LR = 1 × 10<sup>−4</sup>, WD = 10<sup>−5</sup> | 0.9798 | 0.9194 | 0.7671 | 0.7344 | 0.0595 | 0.0932 |
| LR = 1 × 10<sup>−4</sup>, WD = 0 | 0.9796 | 0.9180 | 0.7694 | 0.7371 | 0.0594 | 0.0924 |
| LR = 1 × 10<sup>−4</sup>, WD = 10<sup>−4</sup> | 0.9795 | 0.9173 | 0.7654 | 0.7326 | 0.0597 | 0.0927 |
| LR = 1 × 10<sup>−3</sup>, WD = 10<sup>−5</sup> | 0.9280 | 0.7745 | 0.4389 | 0.3807 | 0.3028 | 0.3751 |
| LR = 1 × 10<sup>−3</sup>, WD = 10<sup>−4</sup> | 0.9273 | 0.7621 | 0.4580 | 0.4029 | 0.2783 | 0.3460 |
| LR = 1 × 10<sup>−3</sup>, WD = 0 | 0.9147 | 0.7502 | 0.4515 | 0.3895 | 0.2661 | 0.3162 |

**Table S6:** Hyperparameter sensitivity comparison between the validation-selected augmentation configuration and its matched standard-shift control. Both configurations use the compact 5CNN+1T architecture and a single training seed (42). The configuration was selected using validation AUPRC; held-out test results are reported only for evaluation. The threshold is selected by maximizing validation F1.

| **Split** | **Configuration** | **AUROC** | **AUPRC** | **F1** | **MCC** | **Threshold** |
|---|---|---|---|---|---|---|
| Validation | RC *p* = 0.5, shift [−8, 8] | 0.9885 | 0.9603 | 0.8915 | 0.8689 | – |
| Validation | RC *p* = 0.5, shift 0 | **0.9898** | **0.9631** | **0.8994** | **0.8782** | – |
| Held-out test | RC *p* = 0.5, shift [−8, 8] | 0.9866 | 0.9414 | 0.8647 | 0.8415 | 0.7340 |
| Held-out test | RC *p* = 0.5, shift 0 | 0.9890 | 0.9485 | 0.8744 | 0.8522 | 0.7271 |

### Supplementary Note 4: Core model and ablations

Supplementary Table S7 reports complete five-seed held-out metrics for MELD, the same-data, same-backbone ISM baseline and each component ablation, which are summarized in Fig. 3A–C.

**Table S7:** Core model and ablation results on the ClinVar standard held-out test set. Results are reported over five seeds as mean ± standard deviation. For each seed, threshold-dependent metrics use the threshold that maximizes validation F1.

| **Model** | **AUROC** | **AUPRC** | **F1** | **MCC** | **Specificity** | **Recall** |
|---|---|---|---|---|---|---|
| MELD | 0.9859 ± 0.0007 | 0.9342 ± 0.0052 | 0.8598 ± 0.0047 | 0.8358 ± 0.0050 | 0.9805 ± 0.0023 | 0.8350 ± 0.0144 |
| Same-backbone ISM baseline | 0.8003 ± 0.0103 | 0.7004 ± 0.0191 | 0.6826 ± 0.0226 | 0.6554 ± 0.0236 | 0.9823 ± 0.0016 | 0.5689 ± 0.0247 |
| No-alpha ablation | 0.9832 ± 0.0006 | 0.9306 ± 0.0014 | 0.8626 ± 0.0033 | 0.8381 ± 0.0032 | 0.9774 ± 0.0025 | 0.8524 ± 0.0155 |
| No-mutation-boost ablation | 0.8929 ± 0.0008 | 0.7938 ± 0.0016 | 0.7610 ± 0.0011 | 0.7423 ± 0.0007 | 0.9908 ± 0.0012 | 0.6452 ± 0.0055 |
| Single-scale ablation | 0.9776 ± 0.0013 | 0.9210 ± 0.0020 | 0.8461 ± 0.0015 | 0.8197 ± 0.0020 | 0.9782 ± 0.0017 | 0.8211 ± 0.0064 |
| No-RC ablation | 0.9696 ± 0.0247 | 0.8880 ± 0.0647 | 0.8082 ± 0.0665 | 0.7758 ± 0.0772 | 0.9728 ± 0.0101 | 0.7806 ± 0.0736 |
| No-delta-regularizer ablation | 0.9756 ± 0.0245 | 0.9097 ± 0.0716 | 0.8320 ± 0.0750 | 0.8024 ± 0.0883 | 0.9731 ± 0.0131 | 0.8178 ± 0.0763 |
| 256-bp context ablation | 0.9734 ± 0.0262 | 0.9011 ± 0.0717 | 0.8263 ± 0.0733 | 0.7956 ± 0.0857 | 0.9717 ± 0.0112 | 0.8143 ± 0.0786 |
| Residual alpha, α<sub>max</sub> = 16 | 0.9857 ± 0.0002 | 0.9383 ± 0.0011 | 0.8607 ± 0.0024 | 0.8365 ± 0.0016 | 0.9791 ± 0.0037 | 0.8423 ± 0.0192 |
| Residual alpha, α<sub>max</sub> = 8 | 0.9856 ± 0.0003 | 0.9382 ± 0.0008 | 0.8611 ± 0.0023 | 0.8369 ± 0.0022 | 0.9791 ± 0.0029 | 0.8430 ± 0.0148 |

### Supplementary Note 5: Shared-subset benchmark for external baselines

For compute-intensive external baselines, a deterministic shared subset of 200k training, 20k validation and 20k held-out test variants was drawn within the original split boundaries. Supplementary Table S8 compares models trained on this subset with the same models trained on full ClinVar, all evaluated on the same 20k test set. Supplementary Fig. S2 plots the comparison against frozen external backbones reported in Table 2, and Supplementary Tables S9 and S10 report the remaining classification, error-rate and calibration metrics for the same models.

**Table S8:** Comparison of MELD and its no-alpha ablation trained on full ClinVar or the shared training subset, evaluated on the same fixed 20,000-variant held-out test set. Values are reported as five-seed mean ± sample standard deviation.

| **Model** | **Training scope** | **AUROC** | **AUPRC** | **F1** | **MCC** | **Recall** | **Specificity** |
|---|---|---|---|---|---|---|---|
| MELD | Full ClinVar | 0.9859 ± 0.0006 | 0.9362 ± 0.0042 | 0.8612 ± 0.0050 | 0.8374 ± 0.0055 | 0.8369 ± 0.0136 | 0.9806 ± 0.0024 |
| MELD | Shared subset | 0.9674 ± 0.0007 | 0.8883 ± 0.0022 | 0.8030 ± 0.0046 | 0.7680 ± 0.0053 | 0.7900 ± 0.0084 | 0.9676 ± 0.0020 |
| No-alpha | Full ClinVar | 0.9831 ± 0.0004 | 0.9269 ± 0.0007 | 0.8633 ± 0.0049 | 0.8391 ± 0.0052 | 0.8527 ± 0.0175 | 0.9777 ± 0.0024 |
| No-alpha | Shared subset | 0.9619 ± 0.0015 | 0.8769 ± 0.0049 | 0.8027 ± 0.0061 | 0.7666 ± 0.0071 | 0.8068 ± 0.0107 | 0.9630 ± 0.0013 |

**Fig S2:**
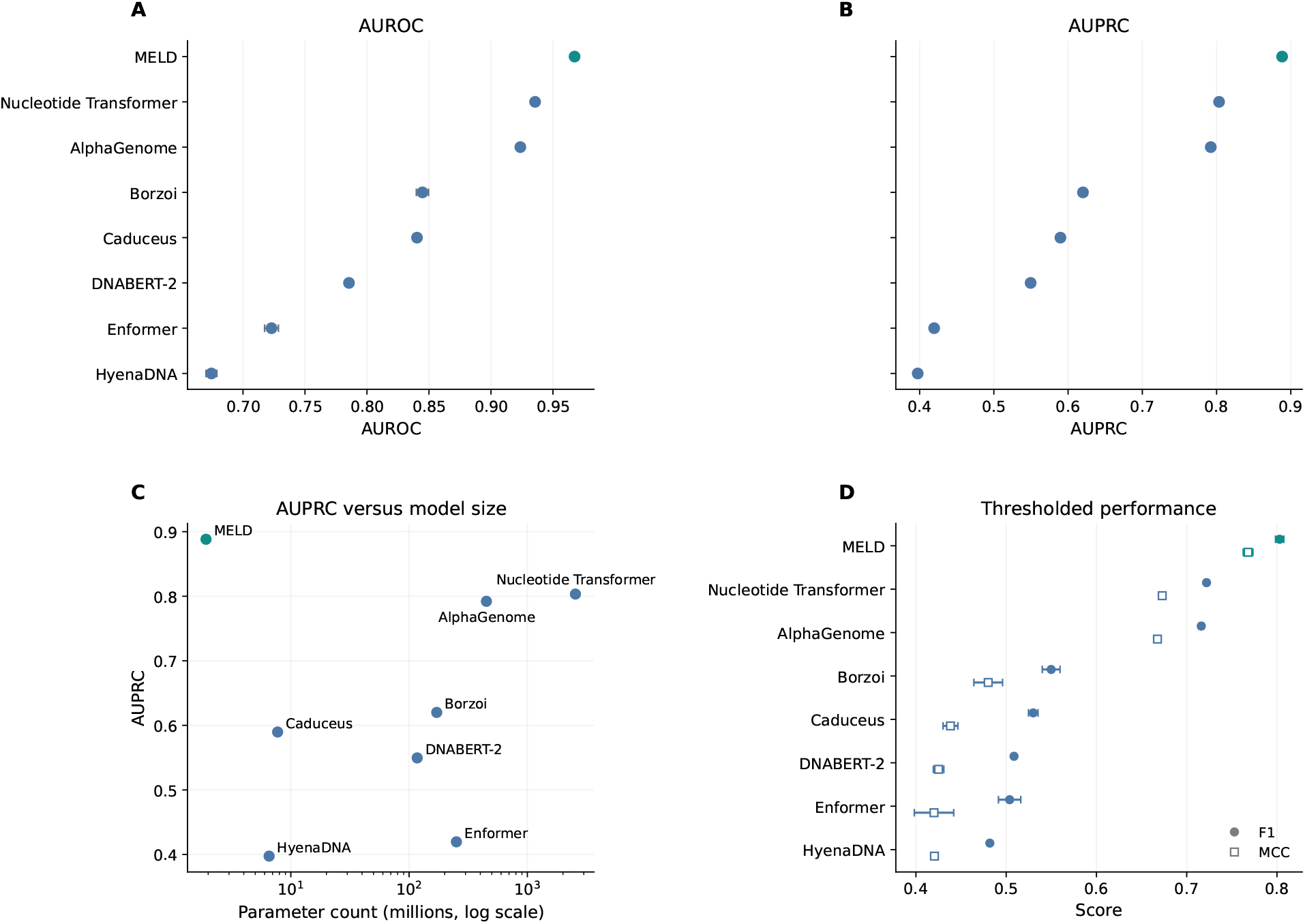
Shared-subset benchmark against frozen external foundation-model baselines. All models were trained on the 200k-variant shared training subset and evaluated on the fixed 20,000-variant held-out test set. MELD (teal) is trained end-to-end; the external backbones (blue) are frozen and only their prediction heads are trained. **(A)** AUROC and **(B)** AUPRC, with models ordered by AUPRC; points and error bars show five-seed mean ± standard deviation. **(C)** Mean AUPRC against parameter count (log scale); counts cover MELD’s complete model and the external backbones excluding their prediction heads. **(D)** F1 (circles) and MCC (squares) at the threshold that maximizes validation F1. Values are listed in Table 2; additional metrics are given in Supplementary Tables S9 and S10.

**Table S9:** Additional classification metrics for the shared-subset external benchmark on the fixed 20,000-variant test set. Values are five-seed mean ± standard deviation, using the same models and thresholds as Table 2. Sequence-to-function backbones provide predicted assay tracks; DNA language models provide trunk embeddings. External backbones are frozen, and only their prediction heads are trained; MELD is trained end-to-end.

| **Model** | **Accuracy** | **Balanced accuracy** | **Precision** | **Recall** | **Specificity** |
|---|---|---|---|---|---|
| MELD | **0.9403 ± 0.0015** | **0.8788 ± 0.0037** | **0.8164 ± 0.0085** | **0.7900 ± 0.0084** | **0.9676 ± 0.0020** |
| Enformer<sup>19</sup> | 0.8537 ± 0.0132 | 0.7013 ± 0.0008 | 0.5320 ± 0.0446 | 0.4812 ± 0.0181 | 0.9214 ± 0.0189 |
| Borzoi<sup>20</sup> | 0.8711 ± 0.0124 | 0.7238 ± 0.0123 | 0.6041 ± 0.0577 | 0.5108 ± 0.0456 | 0.9367 ± 0.0225 |
| AlphaGenome<sup>7</sup> | 0.9159 ± 0.0015 | 0.8231 ± 0.0021 | 0.7456 ± 0.0101 | 0.6890 ± 0.0064 | 0.9572 ± 0.0027 |
| Nucleotide Transformer<sup>21</sup> | 0.9160 ± 0.0016 | 0.8310 ± 0.0054 | 0.7367 ± 0.0145 | 0.7082 ± 0.0152 | 0.9538 ± 0.0045 |
| DNABERT-2<sup>22</sup> | 0.8554 ± 0.0050 | 0.7043 ± 0.0029 | 0.5346 ± 0.0189 | 0.4859 ± 0.0141 | 0.9227 ± 0.0085 |
| Caduceus<sup>23</sup> | 0.8409 ± 0.0092 | 0.7350 ± 0.0046 | 0.4879 ± 0.0241 | 0.5819 ± 0.0235 | 0.8881 ± 0.0151 |
| HyenaDNA<sup>24</sup> | 0.8672 ± 0.0001 | 0.6765 ± 0.0001 | 0.6035 ± 0.0003 | 0.4010 ± 0.0001 | 0.9520 ± 0.0001 |

**Table S10:** Error rates, probability-quality metrics and validation-selected thresholds for the shared-subset external benchmark. Values are five-seed mean ± standard deviation for the models in Table 2. FPR and FNR denote false-positive and false-negative rates, respectively; ECE denotes expected calibration error. Bold marks the best value in each error and calibration column.

| **Model** | **FPR** | **FNR** | **Brier score** | **ECE** | **Threshold** |
|---|---|---|---|---|---|
| MELD | **0.0324 ± 0.0020** | **0.2100 ± 0.0084** | **0.0565 ± 0.0019** | **0.0514 ± 0.0034** | 0.7460 ± 0.0214 |
| Enformer<sup>19</sup> | 0.0786 ± 0.0189 | 0.5188 ± 0.0181 | 0.1701 ± 0.0298 | 0.2348 ± 0.0559 | 0.5805 ± 0.0609 |
| Borzoi<sup>20</sup> | 0.0633 ± 0.0225 | 0.4892 ± 0.0456 | 0.1171 ± 0.0187 | 0.1118 ± 0.0462 | 0.6248 ± 0.0995 |
| AlphaGenome<sup>7</sup> | 0.0428 ± 0.0027 | 0.3110 ± 0.0064 | 0.0932 ± 0.0009 | 0.1327 ± 0.0022 | 0.7029 ± 0.0095 |
| Nucleotide Transformer<sup>21</sup> | 0.0462 ± 0.0045 | 0.2918 ± 0.0152 | 0.0757 ± 0.0025 | 0.0794 ± 0.0100 | 0.6614 ± 0.0169 |
| DNABERT-2<sup>22</sup> | 0.0773 ± 0.0085 | 0.5141 ± 0.0141 | 0.1557 ± 0.0099 | 0.2228 ± 0.0211 | 0.6138 ± 0.0210 |
| Caduceus<sup>23</sup> | 0.1119 ± 0.0151 | 0.4181 ± 0.0235 | 0.1334 ± 0.0020 | 0.1587 ± 0.0056 | 0.5932 ± 0.0278 |
| HyenaDNA<sup>24</sup> | 0.0480 ± 0.0001 | 0.5990 ± 0.0001 | 0.1794 ± 0.0077 | 0.2657 ± 0.0155 | 0.5681 ± 0.0252 |

### Supplementary Note 6: End-to-end fine-tuning of DNA language models

DNABERT-2, Caduceus and HyenaDNA were fine-tuned end-to-end on the full ClinVar training split (1,482,739 variants) and evaluated on the full 54,513-variant held-out test set, the same data used for the MELD reference model. Each backbone was paired with the DeltaEmbedding head used for the frozen baselines, with the encoder learning rate selected on validation AUPRC. Supplementary Table S11 reports all classification and calibration metrics alongside MELD; the headline metrics are summarized in Table 3.

**Table S11:** Held-out test performance of end-to-end fine-tuned DNA language models on the full ClinVar standard benchmark (54,513 test variants), compared with the MELD reference model (Supplementary Table S3). Values are five-seed mean ± standard deviation. Checkpoints were selected using validation AUPRC, and threshold-dependent metrics use the threshold that maximizes validation F1. Bold marks the best value in each row.

| **Metric** | **MELD** | **DNABERT-2 (FT)** | **Caduceus (FT)** | **HyenaDNA (FT)** |
|---|---|---|---|---|
| AUROC | **0.9859** ± **0.0007** | 0.9692 ± 0.0008 | 0.9561 ± 0.0018 | 0.9271 ± 0.0042 |
| AUPRC | **0.9342** ± **0.0052** | 0.9066 ± 0.0006 | 0.8636 ± 0.0041 | 0.7397 ± 0.0129 |
| F1 | **0.8598** ± **0.0047** | 0.8293 ± 0.0024 | 0.7817 ± 0.0035 | 0.7029 ± 0.0074 |
| MCC | **0.8358** ± **0.0050** | 0.8038 ± 0.0025 | 0.7446 ± 0.0033 | 0.6474 ± 0.0086 |
| Balanced accuracy | **0.9078** ± **0.0063** | 0.8784 ± 0.0050 | 0.8610 ± 0.0068 | 0.8315 ± 0.0061 |
| Accuracy | **0.9581** ± **0.0011** | 0.9510 ± 0.0007 | 0.9352 ± 0.0009 | 0.9057 ± 0.0020 |
| Precision | 0.8864 ± 0.0102 | **0.8939** ± **0.0139** | 0.8123 ± 0.0142 | 0.6828 ± 0.0061 |
| Recall | **0.8350** ± **0.0144** | 0.7736 ± 0.0125 | 0.7537 ± 0.0173 | 0.7243 ± 0.0127 |
| Specificity | 0.9805 ± 0.0023 | **0.9832** ± **0.0027** | 0.9682 ± 0.0038 | 0.9388 ± 0.0019 |
| Brier score | 0.0472 ± 0.0009 | **0.0423** ± **0.0009** | 0.0553 ± 0.0017 | 0.0888 ± 0.0041 |
| ECE | 0.0711 ± 0.0014 | **0.0295** ± **0.0022** | 0.0392 ± 0.0062 | 0.0895 ± 0.0102 |

### Supplementary Note 7: Data efficiency

Supplementary Table S12 lists the complete five-seed metrics underlying Fig. 5, obtained with the nested training subsets in Supplementary Table S2 and the update-controlled training protocol described in the main text.

**Table S12:** Data-efficiency comparison of MELD and the same-backbone ISM baseline on the fixed ClinVar held-out test set. Values are reported as five-seed mean ± sample standard deviation. The validation and test sets are identical across all training-set sizes.

| **Model** | **Training size** | **AUROC** | **AUPRC** | **F1** | **MCC** |
|---|---|---|---|---|---|
| MELD | 10k | 0.8090 ± 0.0031 | 0.5448 ± 0.0068 | 0.5020 ± 0.0091 | 0.4140 ± 0.0156 |
| MELD | 50k | 0.9034 ± 0.0024 | 0.7279 ± 0.0080 | 0.6611 ± 0.0103 | 0.6031 ± 0.0135 |
| MELD | 200k | 0.9687 ± 0.0008 | 0.8905 ± 0.0028 | 0.8123 ± 0.0034 | 0.7798 ± 0.0042 |
| MELD | 500k | 0.9820 ± 0.0011 | 0.9250 ± 0.0044 | 0.8471 ± 0.0061 | 0.8206 ± 0.0070 |
| MELD | Full | 0.9861 ± 0.0004 | 0.9367 ± 0.0037 | 0.8619 ± 0.0034 | 0.8379 ± 0.0036 |
| ISM | 10k | 0.5092 ± 0.0103 | 0.2388 ± 0.0112 | 0.2703 ± 0.0048 | 0.0579 ± 0.0855 |
| ISM | 50k | 0.6009 ± 0.0621 | 0.3496 ± 0.0815 | 0.3534 ± 0.0572 | 0.2630 ± 0.0566 |
| ISM | 200k | 0.7698 ± 0.0066 | 0.6425 ± 0.0163 | 0.6148 ± 0.0173 | 0.5679 ± 0.0228 |
| ISM | 500k | 0.7902 ± 0.0087 | 0.6775 ± 0.0135 | 0.6533 ± 0.0169 | 0.6183 ± 0.0210 |
| ISM | Full | 0.7963 ± 0.0048 | 0.6973 ± 0.0049 | 0.6818 ± 0.0058 | 0.6537 ± 0.0076 |

### Supplementary Note 8: Quantitative BRCA1 mutation-effect regression

Supplementary Fig. S3A–C plots the held-out BRCA1 results in Table 5. Supplementary Table S13 and Supplementary Fig. S3D report validation performance of ClinVar-to-BRCA1 transfer as BRCA1-related ClinVar training examples are excluded more strictly. Excluding only exact BRCA1 target alleles gave the strongest validation performance but still allowed ClinVar examples with overlapping local sequence contexts, so the strict window-excluded setting was retained for held-out evaluation.

**Table S13:** Sensitivity analysis of ClinVar-to-BRCA1 transfer under increasingly stringent source-data exclusion. Results are five-seed validation mean ± sample standard deviation.

| **Transfer setting** | **Pearson** | **Spearman** | **RMSE** | **MAE** |
|---|---|---|---|---|
| Random initialization | 0.1573 ± 0.0068 | 0.2144 ± 0.0068 | 1.0245 ± 0.0097 | 0.7799 ± 0.0172 |
| Exact-allele-clean transfer | 0.3735 ± 0.0057 | 0.3843 ± 0.0032 | 0.9634 ± 0.0053 | 0.6901 ± 0.0044 |
| Strict window-excluded transfer | 0.3240 ± 0.0076 | 0.3322 ± 0.0058 | 0.9817 ± 0.0064 | 0.7176 ± 0.0063 |

**Fig S3:**
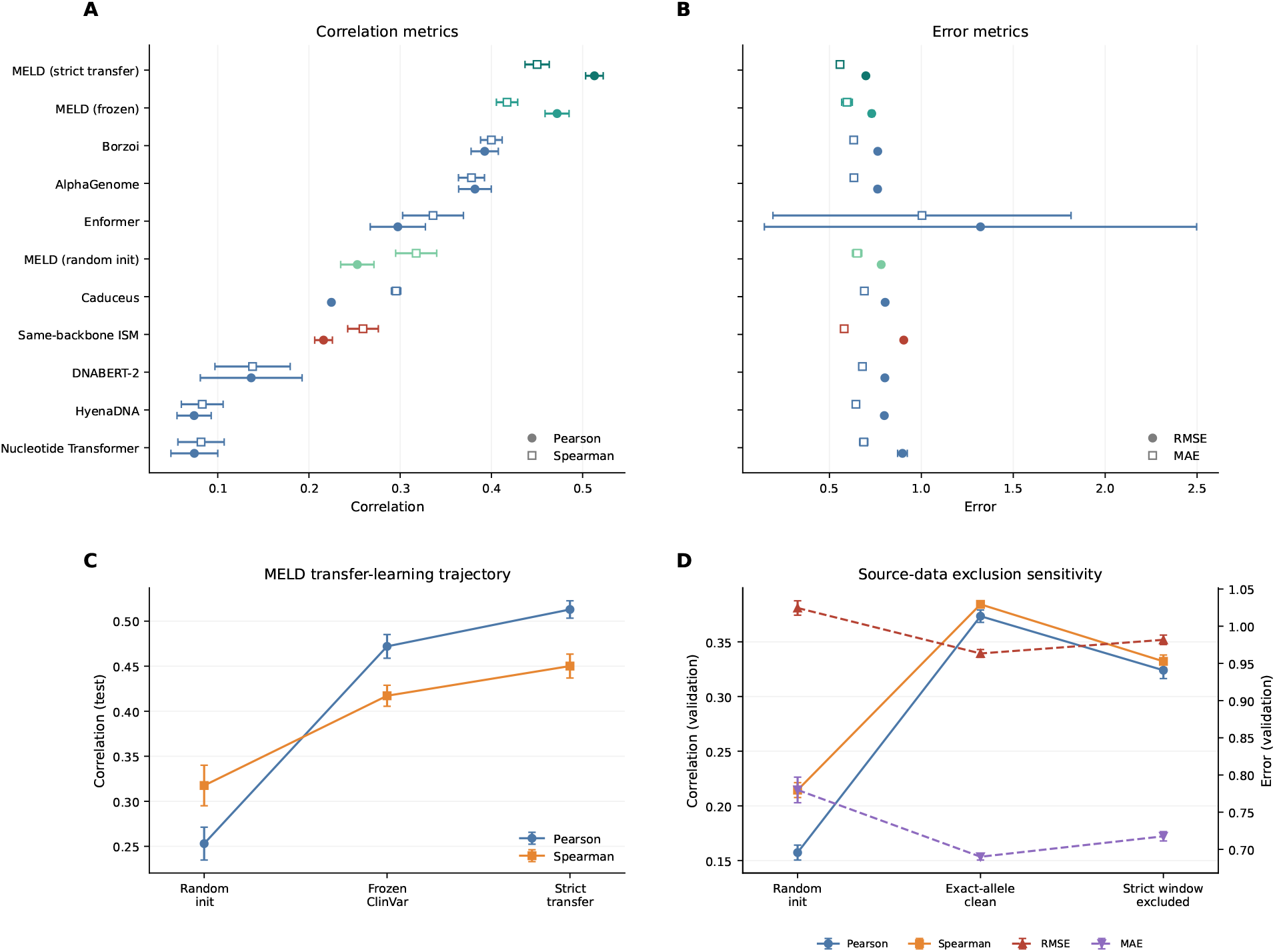
Quantitative BRCA1 mutation-effect prediction and ClinVar transfer. **(A)** Pearson *r* (circles) and Spearman *ρ* (squares) and **(B)** RMSE (circles) and MAE (squares) on the 466-variant held-out BRCA1 test set for MELD after random initialization, with a frozen strict ClinVar-pretrained encoder, and after end-to-end strict ClinVar transfer; for the same-backbone ISM regression baseline; and for seven frozen external encoders. The Enformer error bars in (B) extend beyond the axis (RMSE 1.32 ± 1.18, MAE 1.00 ± 0.81; Table 5). Models are ordered by Spearman *ρ*; points and error bars show five-seed mean ± sample standard deviation. **(C)** Held-out Pearson *r* and Spearman *ρ* of MELD across the three initialization conditions. **(D)** Validation Pearson *r* and Spearman *ρ* (left axis) and RMSE and MAE (right axis) for random initialization, exact-allele-clean transfer and strict window-excluded transfer. Values for (A)-(C) are listed in Table 5 and for (D) in Supplementary Table S13.

